# Tumor-reactive clonotype dynamics underlying clinical response to TIL therapy in melanoma

**DOI:** 10.1101/2023.07.21.544585

**Authors:** Johanna Chiffelle, David Barras, Rémy Pétremand, Angela Orcurto, Sara Bobisse, Marion Arnaud, Aymeric Auger, Blanca Navarro Rodrigo, Eleonora Ghisoni, Christophe Sauvage, Damien Saugy, Alexandra Michel, Baptiste Murgues, Noémie Fahr, Martina Imbimbo, Maria Ochoa de Olza, Sofiya Latifyan, Isaac Crespo, Fabrizio Benedetti, Raphael Genolet, Lise Queiroz, Julien Schmidt, Krisztian Homicsko, Stephan Zimmermann, Olivier Michielin, Michal Bassani-Sternberg, Lana E. Kandalaft, Urania Dafni, Jesus Corria-Osorio, Lionel Trueb, Denarda Dangaj Laniti, Alexandre Harari, George Coukos

## Abstract

The profiles, specificity and dynamics of tumor-specific clonotypes that are associated with clinical response to adoptive cell therapy (ACT) using tumor-infiltrating lymphocytes (TILs) remain unclear. Using single-cell RNA/TCR-sequencing, we tracked TIL clonotypes from baseline tumors to ACT products and post-ACT blood and tumor samples in melanoma patients treated with TIL-ACT. Patients with clinical responses had baseline tumors enriched in tumor-reactive TILs, which were more effectively mobilized upon *in vitro* expansion, yielding products with higher numbers of tumor-specific CD8^+^ cells, which also preferentially infiltrated tumors post-ACT. Conversely, lack of clinical responses was associated with tumors devoid of tumor-reactive resident clonotypes, and with cell products mostly composed of blood-borne clonotypes mainly persisting in blood but not in tumors post-ACT. Upon expansion, tumor-specific TILs lost the specific signatures of states originally exhibited in tumors, including exhaustion, and in responders acquired an intermediate exhausted effector state after tumor engraftment, revealing important functional cell reinvigoration.

## Main

Adoptive cell therapy (ACT) using *in vitro* expanded tumor-infiltrating lymphocytes (TILs) has demonstrated significant efficacy in advanced melanoma^1–3^, and is being tested in other cancer types^4–7^. Although potentially curative in patients who achieve a complete response^1^, the inconsistency of clinical responses to TIL-ACT across patients stresses the need to discern the mechanisms driving therapeutic success^8^. Limitations to therapeutic efficacy have been attributed to date to the scarcity of tumor-specific T cells in tumors of origin^9^, the terminal differentiation of TILs in cell products^2, 10, 11^ or the lack of cell persistence following ACT^12, 13^. Although the absolute frequency of cells recognizing mutational neoantigens is on average low in TIL products^14^, the tumor mutational burden^15^ as well as the frequency of neoantigen-specific clones in TIL infusion products have also been correlated with TIL-ACT efficacy^12, 14^.

Single-cell (sc) sequencing (seq) technologies have revealed transcriptomic signatures of exhausted/dysfunctional states in neoantigen-specific or tumor-specific TIL clonotypes in solid tumors *in situ*^16–22^. Such dysfunctional profiles could limit the ability of tumor-resident TILs to efficiently expand *in vitro* in the context of ACT. Indeed, regardless of the tumor type, a significant loss of clones occurs during TIL expansion *in vitro*, where low frequency clones may outgrow clones with higher *in situ* frequencies^23^. Importantly, cell states in TIL products correlate with persistence in circulation after TIL transfer^13, 24^. However, how TIL repertoires, tumor specificity or original *in situ* cell states affect *in vitro* expansion, *in vivo* blood persistence or tumor engraftment after ACT remain poorly understood. Furthermore, how ACT culture methods impact TIL states, and whether these are effectively reprogrammed upon expansion and subsequent transfer, remain poorly elucidated.

To gain insights into T-cell dynamics and the *in vitro* and *in vivo* fates of cells in the context of TIL-ACT, we interrogated deeply and longitudinally T cells in 13 metastatic melanoma patients who received TIL-ACT in a phase-I clinical trial (NCT03475134). The objective response rate (ORR) in this study was 46.2% (n=6/13)^25^. We examined, by integrated single-cell analyses, the profile of 122’000 T cells, and using bulk TCR-seq we tracked the dynamics of over 2.8×10^6^ clonotypes. We identified profound differences in the original frequency and states of tumor-reactive TILs in baseline tumors, and their dynamic shifts in cell products and in tumors post-transfer in responding and non-responding patients, revealing that clinical responses closely associate with these dynamics.

## Results

### Tumor-resident CD8^+^ vs. blood-borne CD4^+^ clonotype enrichment in cell products and clinical response

The number of adoptively-transferred tumor-specific T cells predicts efficacy in preclinical models and clinical ACT studies^26–28^. For translational purposes, we classified patients as responders (Rs) when they had a complete or partial response (by RECIST criteria v1.1, n=6) vs. non-responders (NRs) when they experienced stable or progressive disease (n=7)^25^. We found that responding patients received more total T cells (**Fig. 1A** and **Supplementary Table 1**), attributed to faster T-cell expansion *ex vivo* (**Extended Data Fig. 1A-B**). Responders’ ACT products (ACTPs) were significantly enriched in CD8^+^ T cells and exhibited higher CD8/CD4 ratios relative to NRs (**Fig. 1B-D** and **Supplementary Table 1**), while CD4^+^ T regulatory cells and γδ T cells, both reputed as immunosuppressive^29, 30^, were enriched in the ACTPs of NRs (**Extended Data Fig. 1C-D** and **Supplementary Table 1**).

**Figure 1.**
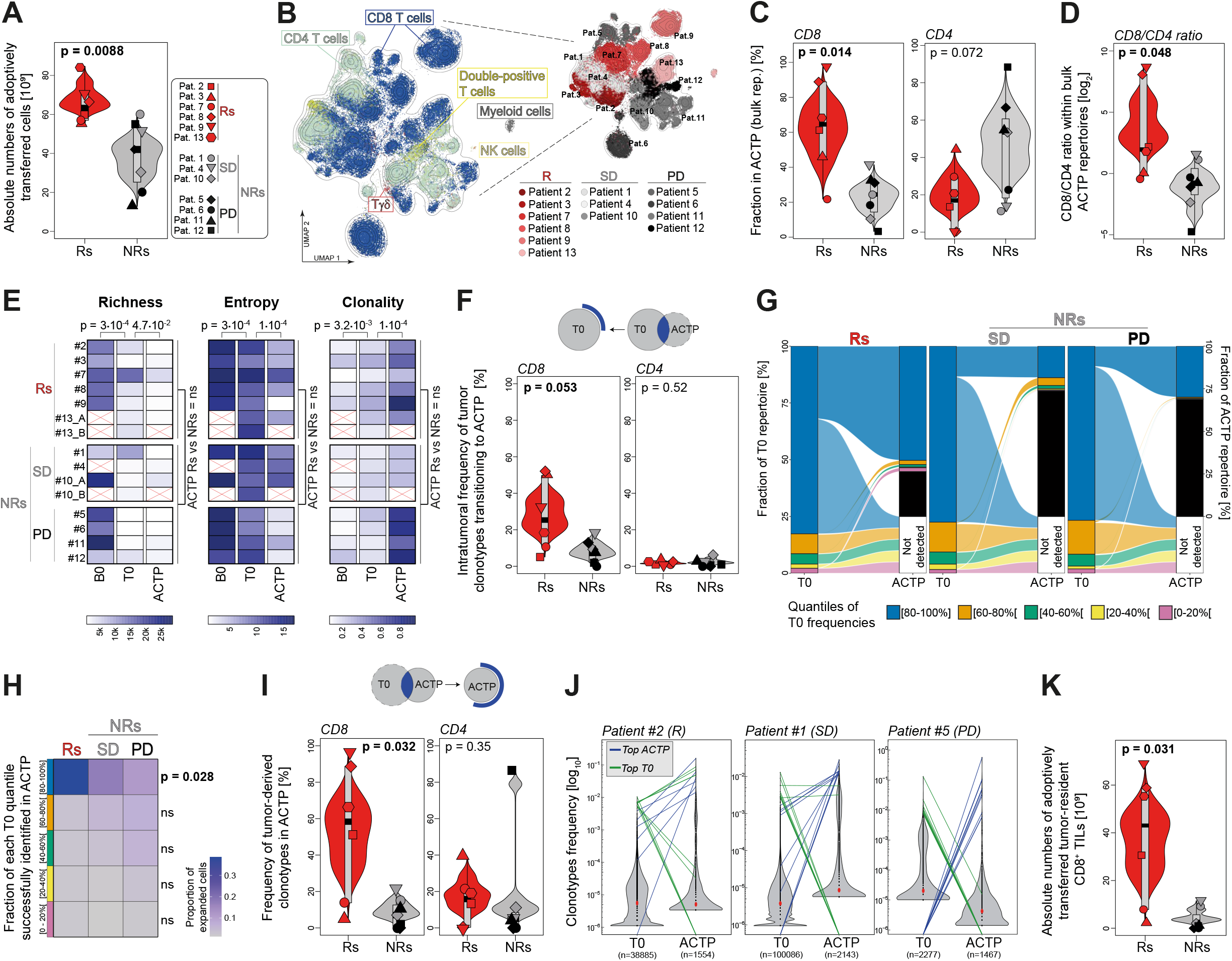
Responders’ and non-responders’ ACT products are enriched in tumor-resident CD8^+^ TILs and bystander CD4^+^ T cells, respectively. **A)** Absolute number of adoptively transferred T cells in responders (Rs) and non-responders (NRs). The different symbols identify the different patients and NRs combine patients with stable disease (SD) and progressive disease (PD). Statistics were performed using a two-tailed t-test. **B)** UMAP projection of ACT product (ACTP) cells from scRNA-seq data of 13 patients highlighting main cell types (left) and patient origin (right). **C-D)** CD8^+^ and CD4^+^ T cell fractions **(C)** and CD8^+^/CD4^+^ T cell ratio **(D)** in Rs’ and NRs’ ACTPs. CD8^+^ and CD4^+^ T cell clones were identified within ACTP bulk TCR repertoires annotated with scRNA-seq. Statistics were performed using two-tailed t-tests. **E)** Heatmap of bulk TCR repertoire metrics showing the richness, Shannon entropy and clonality of baseline blood (B0) and tumor (T0) and ACTP samples. Statistics to compare time points were performed using two-tailed Wilcoxon tests. Comparison between Rs’ and NRs’ ACTP metrics were done using two-tailed Wilcoxon and t-tests. **F)** Intratumoral frequency of CD8^+^ and CD4^+^ T cell clones commonly identified in T0 and ACTP. The intratumoral frequency was determined using T0 and ACTP bulk TCR repertoires annotated with scRNA-seq. Statistics were performed using two-tailed t-tests. **G)** Fractions of tumor repertoires (divided into five color-coded quantiles of equivalent TCR proportions to normalize for clonality) identified in cognate cell products. Intratumoral clones undetectable in ACTP fall in the white box (not detected). ACTP clones undetectable in T0 are represented in black. H) Heatmap showing, for each of the five quantiles of intratumoral clones (from panel G), the proportion of cells successfully expanded (i.e. identified) in ACTP. Statistics between Rs and NRs were performed using two-tailed t-tests. I) Frequency of CD8^+^ and CD4^+^ T cell clones in ACTP commonly identified in T0 and ACTP. The frequency was determined using ACTP and T0 bulk TCR repertoires annotated with scRNA-seq. Statistics were performed using two-tailed Wilcoxon and t-tests. J) Analysis of the top ten clonally expanded clones in T0 (green) and in ACTP (blue) for three representative patients with distinct clinical responses. Violin plots represent the distribution of total bulk TCR repertoires (total number of unique TCRs is indicated in brackets) while colored lines connect clonally expanded TCRs between both compartments. Clones undetectable in a given repertoire are represented below the limit of detection (LOD) of 10^-6^. Cumulative data are shown in **Extended Data Fig. 1F-G. K)** Absolute number of adoptively transferred tumor-resident CD8^+^ T cells in Rs and NRs. Statistics were performed using a two-tailed t-test.

Deep T-cell profiling in tumors at steady state^31, 32^ or in the context of immune checkpoint blockade (ICB) therapy^33, 34^ has revealed an enormous complexity of TCR repertoires. Since T-cell repertoire dynamics have not been investigated to date during TIL-ACT, we sought to interrogate TIL clonotype composition and dynamics in correlation with outcomes. Compared to TILs *in situ* at baseline (T0), we noted that ACTPs showed higher repertoire clonality (i.e. clonal dominance), but lower richness (i.e. fewer distinct clones) and lower Shannon entropy (i.e. lower diversity)^35^, without differences between responders and non-responders (**Fig. 1E**). As expected, circulating T cells collected in blood at baseline (B0) exhibited the highest richness and entropy and the lowest clonality (**Fig. 1E**). We thus asked what fraction of the original tumor-resident clonotypes transitioned into the ACTPs. Interestingly, clonotypes transitioning into ACTPs only represented a minor fraction of tumor-resident CD4^+^ clones, suggesting limited propensity of CD4^+^ TILs to expand under these conditions (**Fig. 1F**). In contrast, tumor-resident CD8^+^ clonotypes dominated ACTPs. Moreover, the fraction of tumor-resident CD8^+^ TIL clonotypes that transitioned into ACTPs was higher in responders than NRs (**Fig. 1F**). In responders, the most dominant (i.e. mostly expanded) tumor-resident clonotypes were likely to transition into the ACTP, unlike lower frequency clonotypes (**Fig. 1G-H**). By contrast, in non-responders, many of the dominant tumor-resident clonotypes failed to mobilize *in vitro* (**Fig. 1G-H**). Accordingly, a larger fraction of the ACTPs, especially the CD8^+^ clones, originated from tumor-resident TILs in Rs (average ∼77%, [46-98%]) compared to NRs (average ∼27%, [0-89%]) (**Fig. 1I** and **Extended Data Fig. 1E**).

We validated these findings by analyzing specifically the top ten dominant clonotypes from each tumor: 7-8 of the top 10 CD8^+^ TIL clonotypes from baseline tumors were identified in the autologous ACTP in responders, while only 2-3/10 did in non-responders on average (**Fig. 1J** and **Extended Data Fig. 1F**). *Vice versa*, 8-9 of the top 10 clonally expanded clonotypes detected in ACTPs originated from baseline tumors in responders, while only 2-3 of the 10 top clones were stemming from tumor in NRs’ ACTPs (**Fig. 1J** and **Extended Data Fig. 1G**). Intrigued by these observations, we surmised that many ACTP clones originated from the circulation in non-responders. Indeed, a large fraction of NRs’ ACTPs, especially the CD4^+^ clones, originated from blood (**Extended Data Fig. 1H**). Consequently, a significantly higher absolute number of tumor-resident CD8^+^ T cells was infused in Rs (**Fig. 1K**).

### Tumor-reactive CD8^+^ clonotypes mobilize in products of responding patients

We surmised that the enrichment in tumor-resident CD8^+^ T cells would translate into increased tumor reactivity in responders’ ACTPs. We used a rapid *in vitro* assay coculturing ACTPs with autologous tumor cells^36^ to quantify tumor-reactive (TR) TILs based on CD137 upregulation (**Fig. 2A**). Responders’ ACTPs contained a significantly higher fraction of CD8^+^CD137^+^ cells (**Fig. 2B**), translating into a significantly higher absolute number of infused tumor-reactive CD8^+^ T cells (**Fig. 2C**), which accurately predicted clinical responses to TIL-ACT (**Fig. 2D**).

**Figure 2.**
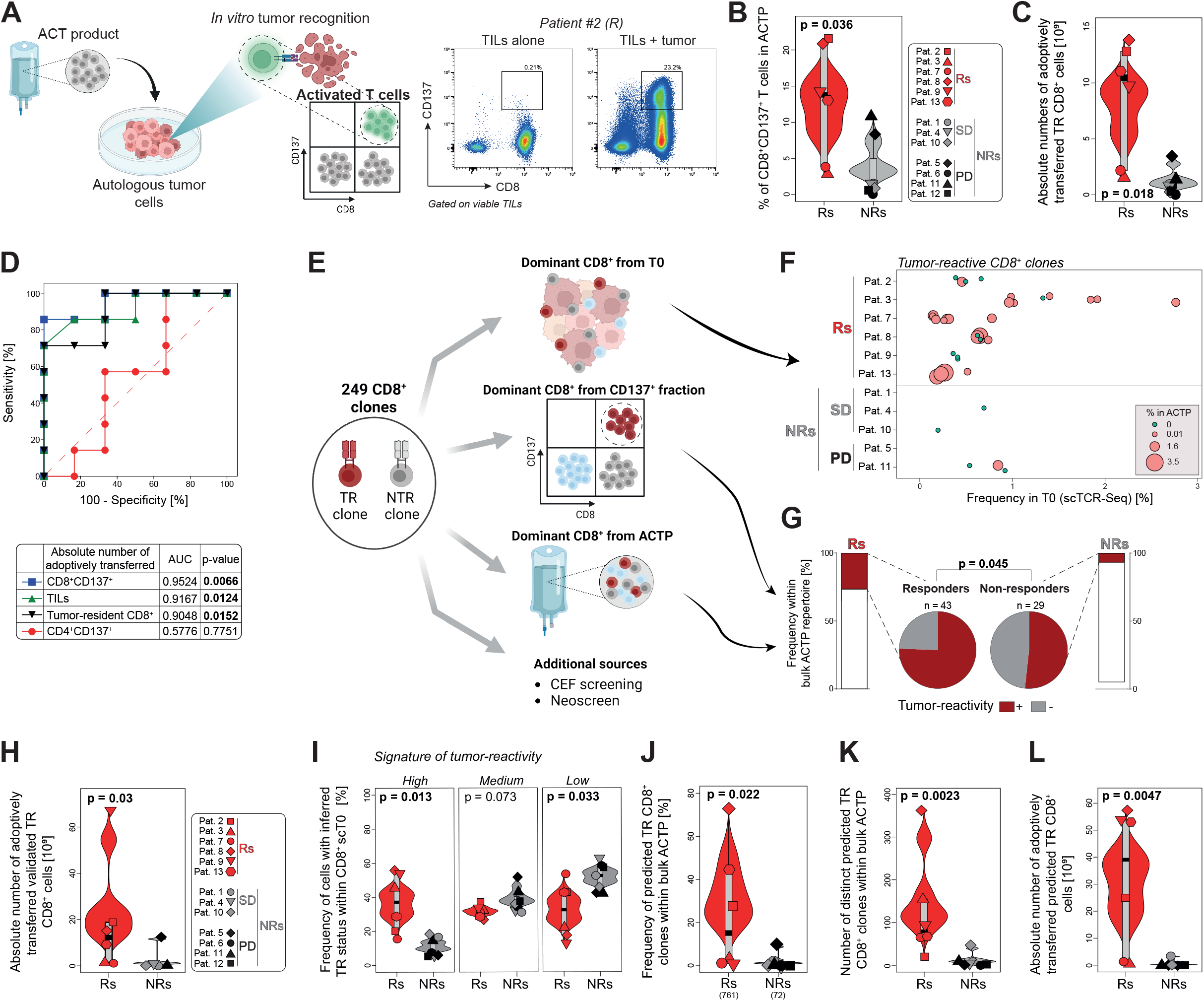
Responders’ tumors are enriched in tumor-reactive clones that successfully expand in ACT products. **A)** Illustration of *in vitro* tumor recognition assay of bulk TILs (from ACTP) and autologous tumor cells (left). Tumor-reactive TILs are defined as those selectively upregulating CD137, as shown in the representative example of flow cytometry data of patient #2 (right). **B**) Frequency of CD137-expressing CD8^+^ T cells after coculture with autologous tumor cells in Rs’ and NRs’ ACTPs. The background levels of CD137 expressed by cognate negative controls (TILs alone) were subtracted. Statistics were performed using a two-tailed t-test. **C**) Absolute number of adoptively transferred tumor-reactive CD8^+^ T cells (i.e. based on CD137 upregulation) in Rs and NRs. Statistics were performed using a two-tailed t-test. **D**) Association between clinical responses and the absolute number of total TILs, tumor-resident CD8^+^ and tumor-reactive (i.e. based on CD137 upregulation) CD8^+^ and CD4^+^ T cells infused per patient. Receiver operating characteristic (ROC) analyses are shown. **E**) Schematic overview of the different origins of individual CD8^+^ clones screened for tumor reactivity. **F**) Individual intratumoral frequencies (in scTCR repertoires) of validated tumor-reactive dominant CD8^+^ T cell clones from baseline tumors shown in **Extended Data Fig. 2A**. Bubble size and color represent clone frequency in ACTP (in scTCR repertoires). **G**) Relative proportion (pies) and corresponding cumulative frequency (histograms) of validated tumor-reactive and non-tumor-reactive CD8^+^ T cell clones (dominant in either bulk or in the CD137-expressing CD8^+^ T cell fractions of ACTP, see methods) in ACTP. Statistics were performed using a Fisher’s exact test. **H**) Absolute number of adoptively transferred tumor-reactive (TR) CD8^+^ T cells (i.e. validated by cloning and screening against autologous tumor cells) in Rs and NRs. Statistics were performed using a two-tailed t-test. **I**) Frequency of all CD8^+^ T cell clones according to their tumor reactivity signature-derived status (*High*, *Mid* and *Low*) in CD8^+^ T cell scTCR repertoires of Rs’ and NRs’ baseline tumors. Statistics were performed using two-tailed t-tests. **J-K**) Frequency (**J**) and diversity (**K**) of predicted tumor-reactive CD8^+^ T cell clones (*High* TR signature status) in ACTP bulk repertoires of Rs and NRs. The total number of tracked clones is indicated in brackets (**J**). Statistics were performed using two-tailed Wilcoxon and t-tests. **L**) Absolute number of adoptively transferred predicted tumor-reactive CD8^+^ T cells in Rs and NRs. Statistics were performed using a two-tailed Wilcoxon test.

Continuing the interrogation specifically of the top clonotypes (**Fig. 2E**), we noted that among the ten dominant CD8^+^ clonotypes found in baseline tumors, a significantly larger fraction was tumor-reactive in responders relative to non-responders (**Extended Data Fig. 2A**, see methods and examples in **Extended Data Fig. 2B-C**). Moreover, a higher fraction of these dominant tumor-resident/tumor-reactive clonotypes transitioned to the ACTPs in responders (average 62.1%, range [0-100%]) than in non-responders (average 11%, range [0-33%]) (**Fig. 2F**). To learn more, we assembled a library of 249 unique CD8^+^ clonotypes from these patients (**Fig. 2E**). These included 72 CD8^+^ TCRs isolated from ACTPs (43 from responders and 29 from NRs), including 53 dominant CD8^+^CD137^+^ clones isolated directly from the *in vitro* tumor reactivity coculture assays, and 19 additional dominant CD8^+^-cell TCRs identified only in ACTPs (example in **Extended Data Fig. 2D**); 98 dominant CD8^+^ clones found in baseline tumors (8 overlapping with the ones found in ACTP); and 87 additional clonotypes that were previously identified as specific to inferred tumor-associated antigens^37^ or viral antigens in these patients. We cloned their αβ-TCR pairs and tested them individually for tumor reactivity (see methods and Supplementary Table 2). A significantly larger proportion of dominant ACTP clones was found to be tumor-reactive in responders, and these clones occupied a higher fraction of the cell products (**Fig. 2G**). Ultimately, responders were infused with a significantly larger absolute number of validated tumor-specific CD8^+^ T cells (**Fig. 2H)**.

Taking advantage of the collection of scRNA/scTCR-seq data from the above validated tumor-reactive CD8^+^ clonotypes, we built a bestowed signature of tumor reactivity for TILs from our melanoma patients progressing after anti-PD1 or anti-PD1/CTLA-4 therapy^25^ (**Supplementary Table 3**). This was concordant with reported signatures of tumor-specific^21^ or neoantigen-specific cells^16–18^ in melanoma or epithelial cancers (**Extended Data Fig. 2E**). Following benchmarking of its ability to predict tumor reactivity (AUC=0.88, p=0.0024, see methods), we used the signature to assign a tumor reactivity score to all CD8^+^ clonotypes identified by scRNA-seq analyses in baseline tumors. We inferred an average of 489 (range [100-1070]) distinct tumor-reactive clonotypes per tumor in responders and 81 (range [7-310]) in non-responders, translating into a significantly higher fraction of the CD8^+^ compartment in responders’ tumors composed of tumor-reactive clones, while NRs’ tumors mostly contained non-tumor-reactive cells (**Fig. 2I**). Using the above bestowed signature, a significantly higher fraction of responders’ ACTPs was composed of cells predicted to be tumor-reactive (**Fig. 2J**), and these included a higher number of distinct clones (**Fig. 2K**). In summary, responding patients received on average 31.8 billion inferred tumor-reactive cells (range [0.7-57.3]) composed on average of 123 unique tumor-reactive clonotypes per product (range [20-361]), while non-responders were estimated to have received on average <1 billion tumor-reactive cells (range [0-3.3]) containing on average only 8 tumor-reactive clonotypes per product (range [0-34], **Fig. 2K-L**). Thus, the tumor-specific payload of TIL-ACTPs can be vastly polyclonal, and clinical response is associated with large numbers of highly polyclonal tumor-specific cells infused. These results provide to our knowledge the first molecular resolution of the TIL-ACT products with respect to tumor specificity and clonality.

### Responders TILs better engraft in tumors post-ACT

TIL persistence in circulation post-ACT is an important predictor of therapeutic efficacy^12, 13^, but how TILs engraft in tumors post-ACT remains to date unexplored. Thirty days post-ACT, we found that tumor repertoires exhibited overall significantly increased clonality, along with lower richness and Shannon entropy, resembling more the autologous ACTPs than the baseline tumors (**Fig. 3A-C**). Similar features were seen in blood 30 days post-ACT (**Extended Data Fig. 3A-C**). In most cases, only a fraction of transferred clonotypes engrafted in tumors (range [14.5-99.6%]) and/or persisted in circulation (range [20.0-99.8%]) (**Fig. 3D** and **Extended Data Fig. 3D**). Such fractions were similar in responders and non-responders and correlated with the overall clonality of ACTPs (**Fig. 3E** and **Extended Data Fig. 3E**). Strikingly, tumor-derived clonotypes from ACTPs effectively infiltrated tumors in patients who experienced objective responses or stable disease, while these cells remained rather confined to the circulation in patients with progressive disease (**Fig. 3F-G** and **Extended Data Fig. 3F-G**). Conversely, blood-borne clonotypes from ACTPs repopulated mostly blood in responders, while they infiltrated tumors in NRs, where blood-borne clonotypes were largely expanded in ACTPs (**Fig. 3F&H** and **Extended Data Fig. 3F&H**).

**Figure 3.**
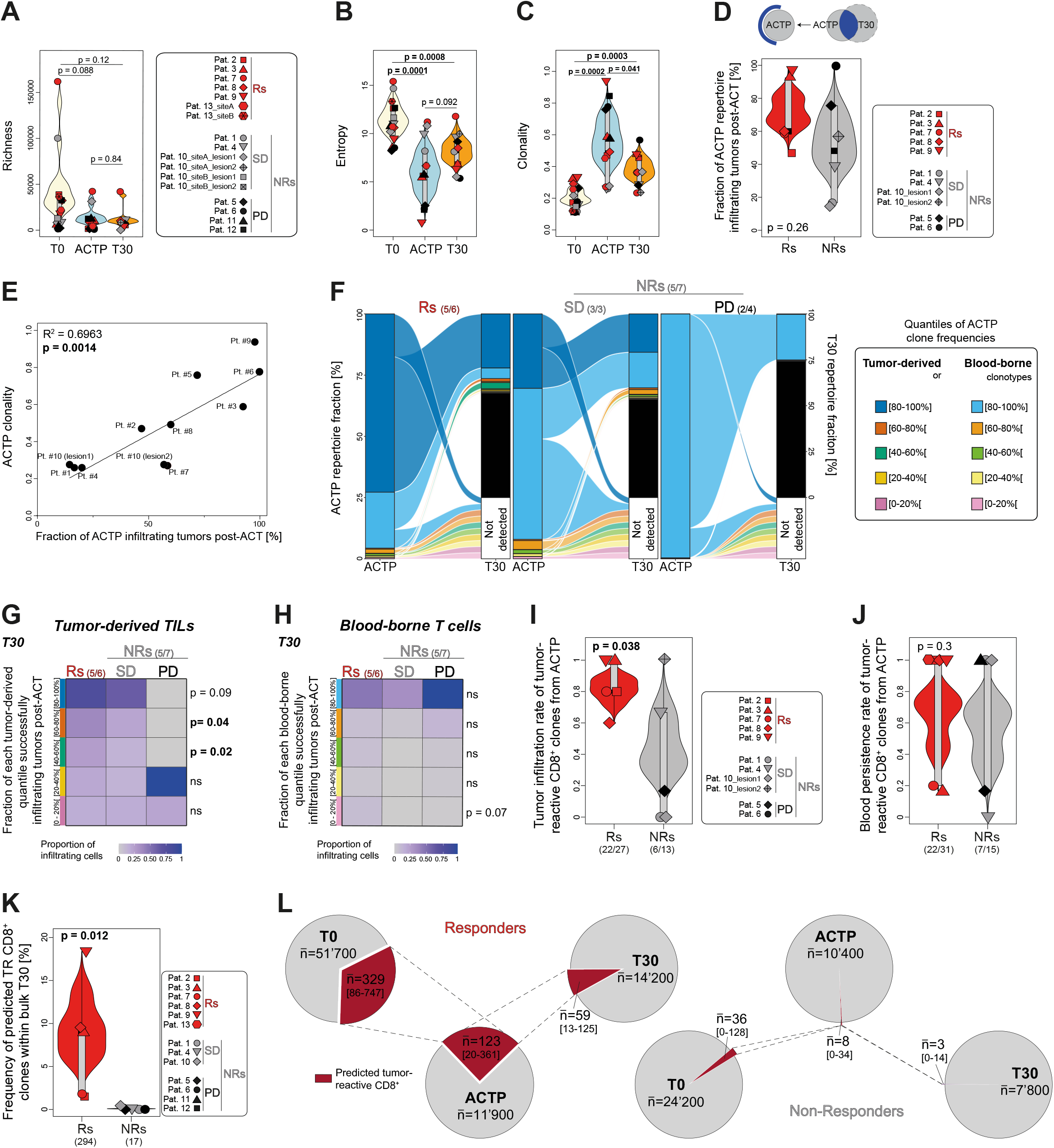
Tumor engraftment post-ACT in responders is associated with tumor-resident TILs from ACT products while it is biased by ACTP repertoire clonality in non-responders. **A-C**) Richness (**A**), Shannon entropy (**B**) and clonality (**C**) comparisons of bulk TCR repertoires between ACTP, pre-ACT (T0) and post-ACT (T30) tumors. Statistics were performed using Wilcoxon and t-tests. **D**) Frequency of clones in ACTP commonly identified in ACTP and T30 in Rs and NRs. The frequency was determined using ACTP and T30 bulk TCR repertoires. Statistics were performed using a two-tailed t-test. **E**) Correlation between ACTP clonality and the fraction of ACTP engrafting tumors post-ACT. Analysis was performed using a linear regression (R^2^ and *p*-value are shown). **F**) Fractions of ACTP repertoires (separated between tumor-derived – detected in T0 – and blood-borne – not detected in T0 but detected in B0 – clones divided into five color-coded quantiles of equivalent TCR proportions to normalize for clonality) identified in cognate post-ACT tumors. ACTP clones undetectable in T30 fall in the white box (not detected). T30 clones undetectable in ACTP are represented in black. **G-H**) Heatmap showing, for each of the five quantiles of ACTP tumor-derived (**G**) or blood-borne (**H**) clones, the proportion of cells successfully engrafted (i.e. identified) in T30. Statistics between Rs and NRs were performed using two-tailed t-tests. **I-J**) Tumor infiltration (**I**) and blood persistence (**J**) rate of validated tumor-reactive CD8^+^ T cell clones dominant in cellular products (and CD137-expressing CD8^+^ fractions, see methods). The total number of infiltrated/persisting clones and the total number of clones identified in ACTP are indicated in brackets. Statistics were performed using one-tailed Wilcoxon and t-tests. **K**) Frequency of predicted tumor-reactive CD8^+^ T cell clones (*High* TR signature status, see methods) in bulk T30 repertoires of Rs and NRs. The total number of tracked clones is indicated in brackets. Statistics were performed using a two-tailed Wilcoxon test. **L**) Pie charts showing the frequency of predicted tumor-reactive CD8^+^ T cell clones (red) in bulk TCR repertoires of pre-(T0) and post-ACT (T30) tumors and cell products (ACTP). The average number of total and predicted tumor-reactive clones, as well as their range, are indicated.

We tracked all dominant ACTP clonotypes that were proven to be tumor-specific. Although persistence in blood post-transfer did not differ between responders and non-responders, we detected a significantly higher rate of tumor engraftment of these cells in responders (**Fig. 3I-J**). We corroborated these findings by tracking all TCRs inferred to be tumor-specific. We found significantly higher numbers of tumor-specific clonotypes engrafting in tumors on day 30 in responders (in total n=294, average of 59) vs. non-responders (in total n=17, average of 3), and detected higher relative frequency of these cells in responders (**Fig. 3K**). Globally, tracking all TCRs from the baseline tumors to day 30 post-ACT, from 329 tumor-reactive clonotypes (among 51,700 bulk TCR sequences) found on average in baseline tumors in responders, 38% transferred into ACTPs, and half of these engrafted in tumors post-ACT (**Fig. 3L**). Strikingly, in non-responders we identified only 36 tumor-resident clonotypes on average that were assigned as tumor-specific, of which only one-fourth transferred into the ACTPs and out of which only one-third engrafted in tumors post-ACT (**Fig. 3L**).

### Exhausted clonotypes preferentially expand *in vitro* and infiltrate tumors post-ACT specifically in responders

Tumor-reactive TILs have been described to acquire mainly exhausted states^21, 22, 38^. Among the 249 CD8^+^ TIL clonotypes characterized, 123 were proven tumor-specific, and these were indeed distributed across canonical exhausted (Tex), precursor-exhausted (Pex) and interferon-stimulated gene (ISG) states, although a large fraction of them were also effector-memory-like (EM-like)^25^ (**Fig. 4A**). As expected, dominant tumor-resident clonotypes that proved to be non-tumor-reactive displayed similar states, although in different proportions (**Fig. 4A**). To extend these learnings, we used the above bestowed signature of tumor reactivity, to interrogate all the baseline CD8^+^ TIL clonotypes (n=10,494) for which we had paired scTCR-seq and scRNA-seq data. We thereby assigned putative tumor specificity to 3,498 clonotypes (see methods and **Extended Data Fig. 4A**) and then examined across all tumor-resident TILs baring those tumor-specific TCRs to identify their state in baseline tumors. As expected, inferred tumor-specific cells were largely distributed in Pex, Tex and ISG states, but again, an important fraction were EM-like, while bystander non-tumor-reactive cells were found in naïve-like and EM-like states (**Fig. 4B**). Reflecting the higher frequency of tumor-reactive CD8^+^ T cells in baseline tumors, we found a higher absolute number of predicted tumor-reactive exhausted cells in responding patients (**Fig. 4C**), translating into a higher proportion of inferred tumor-reactive cells in responders’ Pex and Tex in baseline tumors (**Extended Data Fig. 4B**).

**Figure 4.**
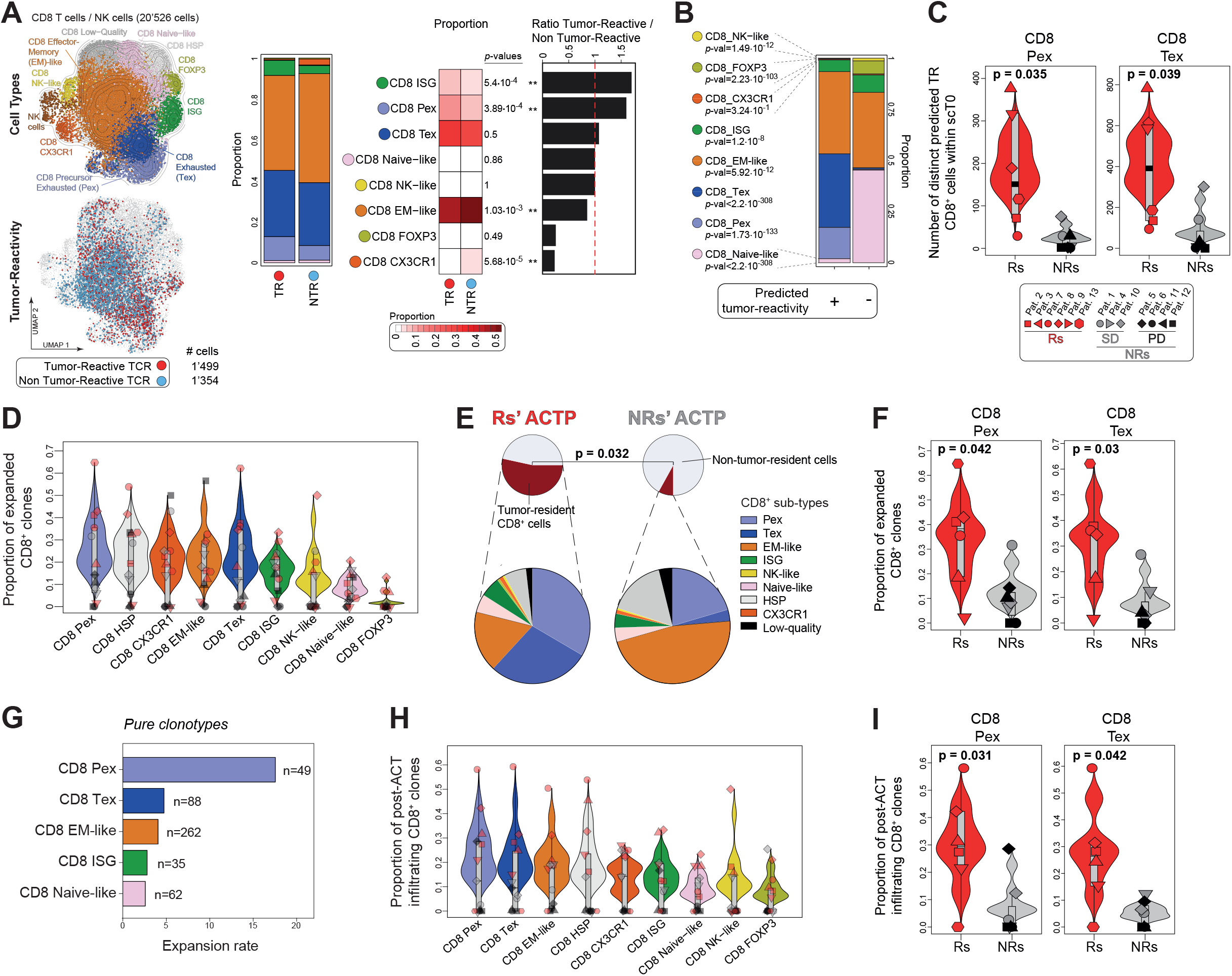
Tumor-reactive cells are enriched in exhausted clusters, which preferentially expand and engraft in tumors post-ACT. **A**) UMAP of CD8^+^ T cells in baseline tumor scRNA-seq data color-coded according to their CD8^+^ T cell subtype (top) and to their validated tumor reactivity status (i.e. validated by cloning and screening against autologous tumor cells) (bottom). The corresponding number of annotated cells is shown below. Distribution of CD8^+^ T cell subtype according to validated tumor reactivity status (middle); heatmap quantifying these proportions. χ^2^-tests were used to compare the proportions of cell types according to the tumor reactivity status and *p*-values were corrected by false discovery rate. The ratio between the number of tumor-reactive and non-tumor-reactive cells within different CD8^+^ T cell subtypes are shown on the right. **B**) Distribution of CD8^+^ T cell subtypes among cells predicted as tumor-reactive (*High* TR signature status, see methods) or non-tumor-reactive (*Low* TR signature status). χ^2^-tests were used to compare the proportions of cell types according to the tumor reactivity status and *p*-values were corrected by false discovery rate. **C**) Absolute number of predicted tumor-reactive (*High* status) cells in intratumoral CD8^+^ Pex and CD8^+^ Tex populations in Rs and NRs. Statistics were performed using two-tailed t-tests. **D**) Proportion of clones from the different intratumoral CD8^+^ T cell subtypes expanding *in vitro* (i.e. detected in cognate bulk ACTP repertoires). **E**) Pie charts showing the contribution of intratumoral CD8^+^ T cell subtypes to the fraction of tumor-resident cells in ACTP. Analyses for each individual CD8^+^ subtype are shown in **Extended Data Fig. 4C**. Statistics were performed using a two-tailed t-test. **F**) Proportion of intratumoral CD8^+^ Pex and CD8^+^ Tex expanding *in vitro* (i.e. detected in bulk ACTP repertoires) in Rs and NRs. Statistics were performed using two-tailed t-tests. **G**) Expansion rate of the *pure* clones, i.e. containing cells found exclusively in a single state (Pex, Tex, EM-like, ISG and naive-like *pure* clones). The number of clones for each population is shown. **H**) Proportion of clones from the different intratumoral CD8^+^ T cell subtypes expanding *in vitro* (i.e. detected in bulk ACTP repertoires) and successfully infiltrating tumors post-ACT (*i.e.* detected in bulk T30 repertoires). **I**) Proportion of intratumoral CD8^+^ Pex and CD8^+^ Tex expanding *in vitro* (i.e. detected in bulk ACTP repertoires) and successfully infiltrating tumors post-ACT (i.e. detected in bulk T30 repertoires) in Rs and NRs. Statistics were performed using two-tailed t-tests.

We next tracked the same TCR clonotypes between ACTPs and tumors. We asked what was the original state of cells that constituted the ACTPs. Expanded CD8^+^ clonotypes were found across different states (**Fig. 4D**). Indeed, a significantly higher fraction of tumor-derived CD8^+^ clonotypes found in the product in responders were originally Pex or Tex, with less EM-like (**Fig. 4E** and **Extended Data Fig. 4C**). Moreover, a significantly higher fraction of clonotypes that were found in exhausted states transitioned in products in responders relative to non-responders (**Fig. 4F**). Strikingly, in non-responders’ ACTPs, tumor-derived clonotypes were EM-like and Pex TILs in the original tumors (**Fig. 4E** and **Extended Data Fig. 4C**). To learn more, we tracked 496 ACTP CD8^+^ clonotypes which could be found exclusively in a single original state in the tumors, i.e. EM-like, Tex, Pex, ISG or naive. Computing the relative expansion of these clones in ACTPs with respect to their original frequency in tumors, we speculated that the frequency of precursor cells giving rise to the product was likely higher among Pex cells than any other TIL subset (**Fig. 4G**). Accordingly, the frequency of tumor-resident Pex *in situ* correlated with the kinetics of TIL expansion and with the absolute number of CD8^+^ T cells in the ACTP and it predicted the absolute number of tumor-reactive CD8^+^ T cells in cell products (**Extended Data Fig. 4D**).

Next, we followed these clones from tumor through the product and in the host post ACT. Remarkably, clones that exhibited an exhausted (i.e. Pex or Tex) state in the tumor of origin efficiently engrafted in tumors post-ACT, but were not always detected in the circulation post-ACT (**Fig**. **4H** and **Extended Data Fig. 4E**). Specifically, we observed significantly higher tumor infiltration by CD8^+^ clonotypes that exhibited Pex and Tex states at baseline in responders than NRs (**Fig. 4I**), whereas there was no association between their detection in blood and response (**Extended Data Fig. 4F**).

### TIL are functionally reinvigorated during *in vitro* expansion

We asked whether *in vitro* expansion somehow reinvigorates tumor-specific CD8^+^ T cells, endowing them with renewed effector features. When examining all cells across all patients, we noted profound changes in the overall transcriptional states upon *in vitro* expansion (**Fig. 5A**). Whereas original tumor-resident CD8^+^ TILs displayed clear distinct canonical states *in situ*, state-specific transcriptional signatures, especially those of canonical Pex and Tex, were altogether attenuated *ex vivo* (**Fig. 5B** and **Extended Data Fig. 5A**). Concurrently, features of effector cells were upregulated in ACTP upon expansion (**Fig. 5B** and **Extended Data Fig. 5A**), in particular cytotoxic genes including granzymes (*GZMA, GZMB, GZMH*), granulysin (*GNLY*), perforin (*PRF1*), *NKG7* and *CX3CR1* (**Fig. 5C**). To shed more light, we tracked all available individual CD8^+^ clones (n=216) identified in single-cell data in all three consecutive samples from each patient: baseline tumors, ACTPs and tumors post-ACT. We confirmed that *in vitro* expanded TILs displayed significantly weaker exhaustion signatures, which was particularly obvious in responders (**Fig. 5D, Extended Data Fig. 5B** and **Supplementary Table 4**), where key markers^22^ including *PDCD1, HAVCR2/*TIM3*, TIGIT, LAG3*, *CXCL13* and *TOX,* the master regulator of T-cell exhaustion, were markedly downregulated relative to the same clones in baseline tumors (**Fig. 5D**). Regulon analyses confirmed that cells underwent important transcriptional reprogramming (**Extended Data Fig. 5B**). Indeed, clonotypes that displayed exhaustion *in situ* transitioned away from this dysfunctional state, with downregulation of EOMES; NR4A1, a key mediator of effector T-cell dysfunction^39^; ZNF831 (zing finger protein 831), which binds to enhancer elements of promoters of several key genes implicated in effector and exhaustion programs; as well as IRF9 and MYOD1, associated with PD-1 signaling, specifically attenuated in responders (**Extended Data Fig. 5B**). Also in responders, we observed upregulation of genes (*MKI67, CDKN2A*) and regulons (MYC and E2F2^40^) associated with cell proliferation (**Fig. 5D** and **Extended Data Fig. 5B**), reflecting the effect of IL-2. Associated with important cell proliferation, the stemness-associated transcription factor (TF) TCF7^41, 42^ was downregulated, including in clonotypes identified as Pex at baseline (**Extended Data Fig. 5B-C**). Key regulons driving effector functions such as FOS and JUN, members of the activator protein 1 (AP-1) family, and RUNX3, a master regulator of effector T-cell gene expression through regulation of T-bet and Eomes^43^, were also downregulated in ACTPs (**Extended Data Fig. 5B**), possibly indicating a state of homeostatic expansion driven by IL-2. Importantly, cytotoxic molecules such as *GNLY* and *GZMB* were upregulated in both responders’ and non-responders’ ACTP cells, while *CD69* and *GZMK,* associated with dysfunctional T cells^24, 44^ were downregulated, revealing increased effector fitness in expanded cells (**Fig. 5D**). Interestingly, responders’ TILs specifically upregulated *CXCR3* upon expansion, a marker associated with tumor infiltration in TIL-ACT^45^ (**Fig. 5D**). Thus, overall, *in vitro* expansion drove a specific cell state, with most notable the elimination of exhaustion programs and upregulation of effector fitness.

**Figure 5.**
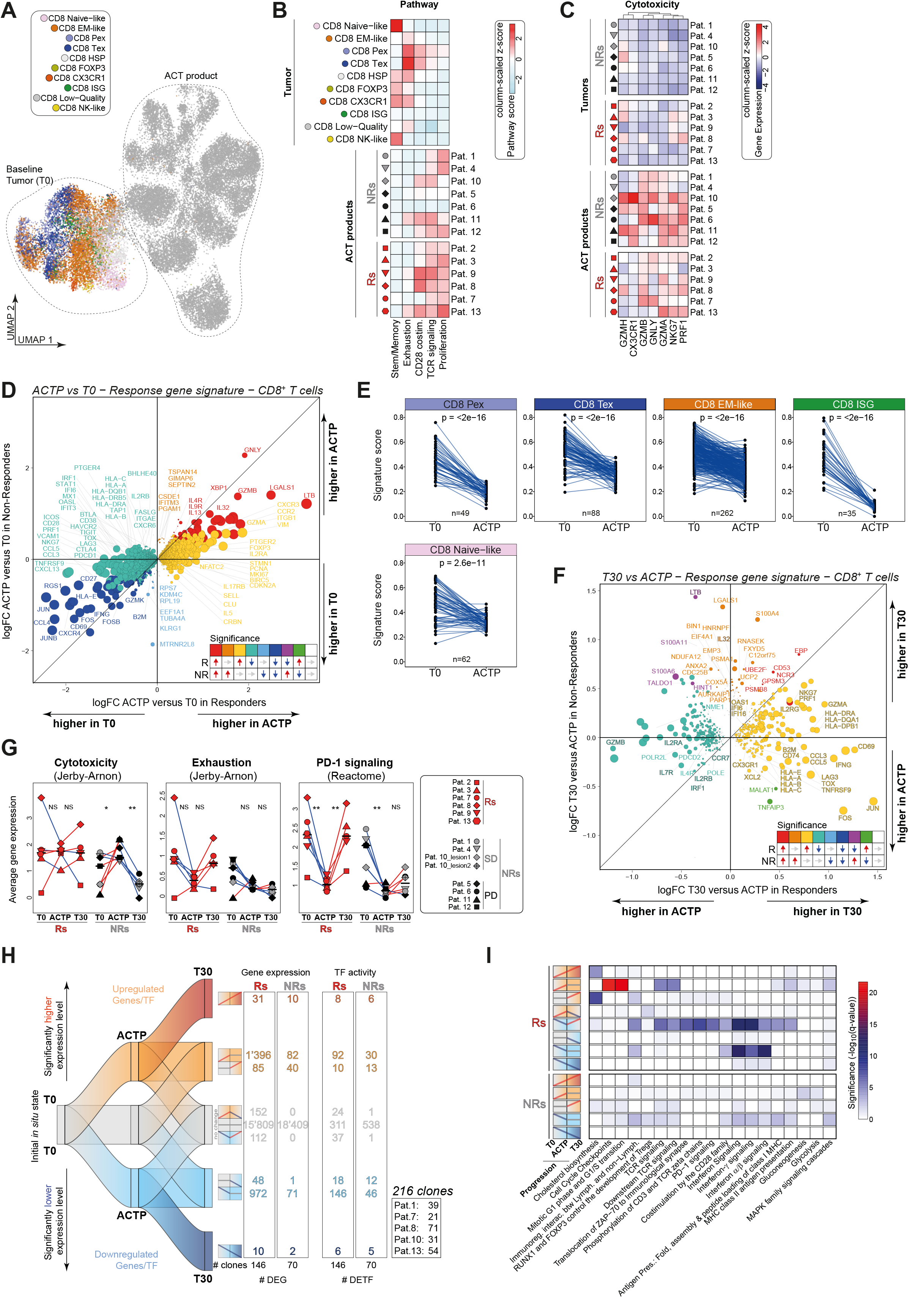
Exhaustion signatures are reprogrammed upon *in vitro* expansion and specifically reacquired in responders’ post-ACT tumor biopsies. **A)** UMAP projection showing clustering of CD8^+^ T cells from baseline tumors (T0) and ACT products of thirteen patients. Cell states from tumors as previously identified (Barras *et al.*) are shown in colors. **B**) Gene signature scores averaged-pseudobulk per CD8^+^ T cell population (for baseline tumors) and per patient (for ACT products). **C**) Cytotoxic genes expression averaged-pseudobulk per patient (for baseline tumors and ACT products). **D**) Scatter plot showing genes differentially expressed in CD8^+^ T cells between ACTP and T0 in Rs (x axis) and NRs (y axis). Colored dots reflect the statistical significance (adjusted *p*-value < 0.05) of each gene in Rs and/or NRs and their directionality (i.e. upregulated or downregulated). **E**) Scoring of gene signatures discriminant for CD8^+^ T cell subtypes (see panel **Extended Data 5A**) in clone-pseudobulk scRNA-seq data (T0 and ACTP) annotated as *pure* (i.e. containing cells found exclusively in a single state at T0). The number of *pure* clones tracked from T0 to ACTP for each CD8^+^ subtype is shown. Statistics were performed using two-tailed paired t-tests. **F**) Scatter plot showing genes differentially expressed in CD8^+^ T cells between T30 and ACTP in Rs (x axis) and NRs (y axis). Colored dots reflect the statistical significance (adjusted *p*-value < 0.05) of each gene in Rs and/or NRs and their directionality (i.e. upregulated or downregulated). **G**) Signature scores (taken from the indicated references) in bulk RNA (T0 and T30) and scRNA-seq averaged-pseudobulk (ACTP) split by clinical response and sample time points. Statistical significance was assessed using paired Student’s t-test. NS: p>0.05; *: p<0.05; **: p<0.01. **H**) Alluvial representation showing the number of genes and transcription factors falling in the nine possible dynamic categories of whether they are significantly up-, down- or not regulated between T0 and ACTP and then to T30. Paired differential gene (DEG) and transcription factor (DETF) analyses were performed on clone average-pseudobulk data and were split according to clinical response. The number of tracked clones is shown on the right. **I**) Reactome pathway analysis showing selected pathways that are the most significantly representing the genes falling in the dynamic categories described in panel **H**.

To test that such reprogramming could be observed also in cells originally situated only in the exhausted compartment *in situ*, we tracked specifically the 496 CD8^+^ clonotypes found originally in one cell state only, as above (see **Fig. 4G)**. We confirmed that the IL-2 induced state also occurred in cells that originally exhibited uniquely an exhausted, ISG or EM-like state (**Fig. 5E)**, indicating that cell reinvigoration upon expansion could affect both exhausted and EM-like cells.

### TIL exhaustion is reacquired post-transfer in responding tumors

Since reprogrammed tumor-resident TILs from responders preferentially infiltrated tumors post-ACT, we asked how these cells adapted to antigen encounter and chronic stimulation *in vivo*. Uniquely in responders’ TILs, we noted a significant upregulation post-ACT of genes implicated in anti-tumor immunity, reflecting tissue-residency (e.g. *CD69*^46^) and effector function (*TNFRSF9*, *GZMA*, HLA-class II molecules, *IFNG*, *CCL3*, *CCL5* and *PRF1* as well as *FOS* and *JUN*), along with exhaustion markers (*LAG3*, *TOX)*, while TCF7 was downregulated (**Fig. 5F, Extended Data Fig. 5D** and **Supplementary Table 4**). Inferred regulon analysis supported important reprogramming of cells towards effector (FOS, JUN, RUNX3) and exhaustion state post-ACT (TBX21, NR4A1 and EOMES), with interferon activation (IRF2, IRF5 and IRF9) in responders (**Extended Data Fig. 5D**). By contrast, this functional adaptation did not occur in non-responders, where few genes not directly linked with an immune response were upregulated (**Fig. 5F**).

These findings indicate that specifically in responders, relevant tumor-specific CD8^+^ TILs shed exhaustion upon expansion but re-acquire a state of antigen-experienced effector cells post-ACT in tumors. To learn more, we analyzed bulk or pseudo-bulk RNA-seq data, which revealed that – specifically in responding patients – cells regained a gene signature of PD-1 signaling, heralding active immune recognition post-ACT, but simultaneously retained the specific cytotoxicity signature acquired during IL-2 expansion. This adaptation *in vivo* was not observed in non-responders (**Fig. 5G**).

We tracked 216 CD8^+^ clonotypes from the original tumors through expansion in ACTPs and then infiltration in tumors post-ACT to investigate these dynamics in more detail. We confirmed their transcriptional programming away from exhaustion during *in vitro* expansion. In responders we validated that important preservation of transcriptional states acquired during expansion, including proliferation and effector features, were retained post-transfer, while an attenuated state of exhaustion was acquired (**Fig. 5H, Extended Data Fig. 6A** and **Supplementary Table 4**). Reactome analysis confirmed PD-1 signaling downregulation upon *in vitro* expansion and re-upregulation post-ACT in tumors specifically in responders, along with TCR and interferon signaling (**Fig. 5I**). Moreover, in responders, cells also exhibited upregulation of cholesterol biosynthesis and MAPK signaling programs post-ACT, implying effector-cell activation (**Fig. 5I**). Interestingly, *TOX* was lost upon expansion and re-upregulated in tumors by day 30, albeit at lower levels than baseline tumors (**Extended Data Fig. 6A**). Importantly, *TNFRSF9*, *PRF1*, *IFNG*, *CD69* were also restored in responders, while *GNLY* and *GZMA* were higher than baseline tumors. These results confirm that TIL-ACT indeed drove important cell reprogramming, resulting in functionally reinvigorated effector cells that although acquired TOX and exhaustion features in tumors post-ACT, were also retaining activated effector-cell features and capable of driving objective tumor responses in patients who failed prior PD-1 therapy.

## Discussion

The absolute number of infused cells is critical for the success of CAR-T therapy^27, 28^, and TIL-based therapy is no exception^11^. In this study we provide unprecedented high-resolution analysis of the cell infusion products during TIL-ACT in melanoma patients. Unsurprisingly, responses were associated with a higher number of tumor-reactive cells infused, the vast majority of which were CD8^+^. Patients who responded exhibited a higher frequency of tumor-reactive cells in tumors at baseline. Moreover, a higher fraction of dominant tumor-resident and tumor-reactive CD8^+^ clonotypes transitioned into the product in responding patients relative to non-responding ones. Strikingly, non-responders’ ACTPs were rather overpopulated by CD4^+^ clones, largely deriving from circulating cells. The specific antigens recognized by TILs remain to date elusive. Indeed, the frequency of cells recognizing canonical shared tumor-associated antigens such as cancer testis antigens or epitopes derived from overexpressed lineage antigens is minimal in TIL products^47^. Furthermore, the average occurrence of lymphocytes recognizing mutational neoantigens is also low^14^. Thus, many of the TIL clonotypes that presumably drive tumor responses remain to date orphan, i.e. without an identified cognate antigen. Nevertheless, our study reveals the extraordinary repertoire richness of tumor-specific cells delivered through TIL-ACT. Using a bestowed gene signature revealing tumor reactivity *in situ*, we tracked cells into the products, and inferred that responding patients received a highly polyvalent cell product containing on average ∼30-fold more tumor-reactive cells in total and ∼15-fold higher number of distinct clonotypes relative to non-responders. A maximum number of 361 distinct tumor-reactive clonotypes were infused, illustrating the potential of TIL-ACT.

Strikingly, even in responders, a rather small fraction of all tumor-resident clonotypes expanded successfully, and ∼60% of inferred tumor-reactive clonotypes failed to reach the end of expansion, in spite of being highly abundant in the original tumors. Further attrition was observed upon transfer, where only half of the infused tumor-reactive clonotypes finally engrafted in tumors in responding patients. In non-responding patients the loss of relevant clones was even more detrimental, leading to products with insufficient mass of tumor-reactive cells and to post-ACT engraftment of irrelevant blood-derived clonotypes in tumors. Elucidation of the underlying mechanisms that enable successful cell expansion and persistence will help enhance the efficacy of TIL-ACT.

The quality and functional characteristics of T cells play a key role in determining the success of ACT. Persistence of transferred cells is a critical parameter associated with durable clinical responses to ACT^12, 13, 48, 49^, and cells with stem-like characteristics provide a powerful therapeutic product even when administered in small numbers^50^. In fact, the presence of CD39^-^ CD69^-^ memory-progenitor CD8^+^ cells capable of self-renewal, expansion and persistence, which included neoantigen-specific cells, was associated with complete cancer regression and TIL persistence^24^. Here we found that infused cells displayed globally unique transcriptional programs associated with reprogramming likely related to the homeostatic expansion by IL-2. *In vitro* expanded tumor-specific cells lost prior gene signatures, notably exhaustion, while maintaining effector functions characterized by CD28 costimulation, TCR signaling and upregulation of cytotoxic genes, thus overall acquiring a reinvigorated functional state. Response to ICB occurs in tumors with pre-existing clonally expanded effector or PD-1^high^ T cells within tumors^33, 51^, and durable responses rely on the activation of precursor-exhausted T cells in preclinical and human melanoma patients^52, 53^. Importantly, in our study we included only patients who had failed prior ICB, which may have biased the departing state and/or impacted the ability of precursor cells to maintain stemness upon IL-2 expansion.

As shown in our study and in prior larger studies, TIL-ACT can drive tumor regression in tumors that failed immune checkpoint blockade^3, 54^. Indeed, a subset of patients failing ICB harbor meaningful frequencies of tumor-reactive TILs, which fail to reinvigorate properly with ICB, yet they are properly invigorated by TIL-ACT. In our investigation, tumor-reactive clonotypes that effectively expanded were distributed in different states in the tumors of origin, including Tex, Pex, ISG and EM-like. To shed light on individual cell fates, we tracked tumor-specific clonotypes from the product back to the original tumors, and found that for many successfully expanded clonotypes we could identify cells among different canonical states, lending support to the notion that expanded tumor-specific TILs could indeed derive from both memory-like and exhausted precursors^25^. Given their precursor stem-like properties, Pex cells are the most likely candidates to expand in response to IL-2, corroborated by analysis of individual clonotypes with only one identifiable *in situ* state. However, we have previously described also a subset of Tex cells embedded in intraepithelial tumor myeloid niches where they can receive CD28 costimulation by activated tumor-resident dendritic cells^38^. We have shown that this subset of tumor-reactive TILs with canonical Tex features but expressing CD28 in fact also express the IL-2 receptors and can sense IL-2 *in situ*, judged by detection of pSTAT5 and proliferation by mass cytometry^38^. This subset of exhausted CD8^+^ TILs was also identified in cell doublets with DCs in this melanoma cohort analyzed here^25^, indicating that a subset of Tex cells could also expand in response to IL-2. STAT5, a key downstream mediator of the IL-2 programs, was recently shown to rewire exhaustion, downregulating the master transcriptional regulator TOX and imparting an intermediate exhaustion state endowed with stronger effector features in mice^55^. Thus, IL-2 could effectively reprogram TILs and attenuate prior state commitments, including exhaustion, driving cells to enter a common state of reinvigorated effector cells. Maintaining and enhancing stemness properties during such cell reprogramming upon expansion should be an important goal of TIL-ACT development.

Tumor-reactive cells expanding out of tumor-resident clonotypes were prone to infiltrate tumors more efficiently, likely related to the expression of homing receptors such as CXCR3, a critical mediator of intratumoral T-cell homing^45, 56^. Tumor homing of tumor-specific clones also depends on TCR affinity^45^, indicating that a pool of tumor-specific TILs with high affinity TCRs were included in the products and infiltrated tumors of patients who benefitted from TIL-ACT. Tumor-specific clones that successfully engrafted in tumors re-acquired features of TOX^+^ exhaustion post-ACT, albeit exhaustion marker and TOX expression was attenuated relative to baseline, and cells expressed higher levels of proliferation and effector markers, resembling phenotypically mouse effector-like TOX^int^ T cells^57^, supported also by the upregulation of T-bet reacquired in tumors post-ACT.

Thus, IL-2 reinvigorates tumor-specific clonotypes, positioning cells in a superior effector state with attenuated exhaustion which persists post-ACT, explaining the success of TIL-ACT even after ICB failures. Although such effector-exhausted intermediate cells have been linked to cell reinvigoration and efficacy of ICB in mice^57^, here we show for the first time that this reinvigorated state is associated with clinical response to TIL-ACT in patients. Furthermore, these TOX^int^ cells could be best positioned to benefit from ICB, providing a strong rationale for combining ICB with TIL-ACT. Cell engineering can provide elegant solutions to endow TILs with the ability to counter PD-1 inhibition^58^ or prevent them from reacquiring exhaustion post-transfer^55, 59^. This study provides novel mechanistic insights linking TIL clonal specificity and state dynamics with ACT efficacy, opening the door to biomarker development for patient selection as well as novel hypotheses for improving TIL-based therapies.

## Methods

### Patients and study design

Patients were enrolled in a single-center phase I clinical study of TIL therapy (ClinicalTrials.gov NCT03475134). This trial was approved by the ethics committee of Canton de Vaud and was conducted in accordance with the principles of Good Clinical Practice, the provisions of the Declaration of Helsinki, the ICH, as well as all national legal and regulatory requirements. All patients provided written informed consent. The characteristics and treatments of the thirteen eligible patients with metastatic melanoma constituting the “per protocol” cohort of the study as well as the study design and conduct are extensively described in Barras *et al.*^25^. Briefly, eligible patients were adults with histologically proven unresectable locally advanced (stage IIIc) or metastatic (stage IV) cutaneous melanoma who have progressed on at least 1 standard first line therapy, including but not limited to chemotherapy, BRAF and MEK inhibitors, anti-CTLA4, anti-PD-1 or anti-LAG3 antibodies and/or the combination. TILs were expanded *in vitro* from patients’ tumor deposits resected by surgery under high doses of interleukin-2 (IL-2)^60^. The patients received a lymphodepletion regimen consisting of fludarabine (25 mg·m^-2^ per day) for 5 days and cyclophosphamide (60 mg·kg^-1^ per day) for 2 (overlapping) days, followed by a single infusion of TILs with concomitant and follow-up intravenous boluses of high dose IL-2. A blood sample was obtained at the same time as the tumor surgery and 30 days post-ACT. Peripheral blood mononuclear cells (PBMCs) were then cryopreserved and stored in a liquid nitrogen container. In addition, an accessible tumor site was biopsied 30 days post-ACT, and one biopsy was snap frozen (for bulk TCR-seq) whereas a second one (if available) was cryopreserved if not freshly used (for single-cell analysis). We observed an objective response rate (by RECIST v.1.1) of 46.2% (6/13 patients - 95% C.I.: 19.2% – 74.9%). Two patients (15.4%) with complete response (CR) and four (30.8%) with partial response (PR) were recorded and classified as responders (Rs). For translational purposes, we classified four patients with progressive disease (PD) and three patients with stable disease (SD) as non-responders (NR, n=7), with two having PD recorded at the next scan while one had durable SD up to 10 months (see Figure 1, Barras *et al.*^25^). Treatment resulted in a median progression-free survival (PFS) of 5.6 months (95% CI 1.2-8.5), and a median overall survival (OS) of 8.8 months (95% CI 7.5 – not reached, with a median follow-up of 45.4 months, IQR: 40.9 – 50.3).

### Sample processing

Resected tumors were minced into 1-2 mm^2^ pieces and, along with available post-infusion biopsies, cryopreserved in 90% human serum + 10% dimethyl sulfoxide (DMSO) and additional pieces were snap frozen for bulk RNA extraction. For single-cell experiments, both frozen and fresh material were used as starting material. PBMCs were isolated from blood collected in EDTA tubes and cryopreserved in 90% human serum + 10% DMSO. TILs were generated in the GMP manufacturing facilities of the Centre of Thérapies Expérimentales (CTE) of the Centre Hospitalier Universitaire Vaudois (CHUV) under specific conditions previously described^60^. Media changes and cell counts were performed on specific days during the pre-REP and REP phases. REP harvest was performed on day 14 and the final product bag was sealed until transport to the clinic. QC sampling was performed to assess various parameters, including final cell count listed in Barras *et al.*^25^. Some TILs of the infusion product were cryopreserved in 90% fetal bovine serum + 10% DMSO for translational studies. A summary of sample availability for the different technologies used is shown here.

|  | B0 | T0 |  | ACTP |  | B30 | T30 |  |
| --- | --- | --- | --- | --- | --- | --- | --- | --- |
| Patient | Bulk TCR-Seq | Bulk TCR-Seq | Sc RNA/TCR-Seq | Bulk TCR-Seq | Sc RNA/TCR-Seq | Bulk TCR-Seq | Bulk TCR-Seq | Sc RNA/TCR-Seq |
| #1 | X | X | X | X | X | X | X | X |
| #2 | X | X | X | X | X | X | X | - |
| #3 | X | X | X | X | X | X | X | - |
| #4 | X | X | X | X | X | X | X | - |
| #5 | X | X | X | X | X | X | X | X |
| #6 | X | X | X | X | X | X | X | X |
| #7 | X | X | X | X | X | X | X | X |
| #8 | X | X | X | X | X | X | X | X |
| #9 | X | X | X | X | X | X | X | - |
| #10 | X | X * | X * | X | X | X | X ** | X ** |
| #11 | X | X | X | X | X | X | - | - |
| #12 | X | X | X | X | X | X | - | - |
| #13 | X | X * | X * | X | X | X | - | X |

Blood and tumor at baseline (B0/T0), ACT product (ACTP) and blood and tumor 30-days post-ACT (B30/T30)
\* Two tumor sites were surgically removed. Samples were analyzed separately for bulk and pooled for single-cell analyses. For patient #10, two pre-REP were launched and pooled into one single REP resulting in one ACT product.
\*\* Two tumor biopsies were obtained, lesion 1 and 2. For bulk analysis, both lesions were available, while for single-cell, only lesion 1 could be processed.

### Generation of tumor cell lines

Tumor cell lines were established from fresh tumor fragments digested in a mix of collagenase type I (Sigma-Aldrich) and DNAse (F. Hoffmann-La Roche) in R10 medium (RPMI 1640 GlutaMAX (Gibco) supplemented with 10% fetal bovine serum (FBS) (Gibco), 100 IU·mL^-1^ of penicillin, 100 μg·mL^-1^ of streptomycin (Bio-Concept)) at 37°C for 90 min. The cell suspension was then harvested and passed through a cell strainer to remove potential remaining tissue pieces. Cells were plated in R10 and incubated at 37°C with 5% CO_2_. Culture medium was replenished every 2-3 days and cultures were split when reaching 80% confluence. To this end, tumor cells were gently detached with Accutase (ThermoFisher Scientific), split and R10 medium was fully replenished. The day before any coculture assay with bulk TILs or for TCR cloning (tumor reactivity described below), tumor cells were incubated 24 hours in R10 medium (i.e. supplemented or not with 200 ng·mL^-1^ IFNγ (Miltenyi Biotec)). Tumor cell lines could be generated for all patients, except patient #6 and #12, for whom the product’s tumor reactivity assay of the product was conducted using tumor digest and no TCR screening was carried out.

### Flow cytometry panel on ACT products

A fraction of TILs from ACT product was freshly received and, after washing, cells were stained for 20-30 min at 4°C with an antibody mix composed of: Zombie UV viability dye (BioLegend), anti-CD45RA (Beckman), anti-CD3 (BD Bioscience), anti-CD27 (eBioscience), anti-CD45 (BioLegend), anti-CD4 (BioLegend), anti-CD8 (BioLegend), anti-PD1 (BioLegend), anti-TIM-3 (R&D), anti-41BB/CD137 (BioLegend), anti-CD56 (BioLegend), anti-CCR7 (BioLegend), anti-CD25 (BioLegend) and anti-CD127 (BioLegend). After a washing step, samples were finally diluted in 500 µL of PBS and analyzed entirely on a BD LSRFortessa Cell Analyzer (BD Biosciences). Data were then analyzed using FlowJo software (LLC). The fractions of CD4^+^ and CD8^+^ T cells were then used for all the downstream analyses where total number of cells infused in patients are highlighted, except when it was inferred from bulk TCR repertoire overlaps annotated with scRNA-seq data (indicated in the legends).

### Isolation of tumor-reactive fraction of ACT products

To assess antitumor reactivity of TILs from ACT products, TILs were thawed and allowed to recover for 48 hours in R8 medium (RPMI 1640 GlutaMAX (Gibco) complemented with 8% human AB serum (Bio West), non-essential amino acids, 100 mM HEPES, 1 mM sodium pyruvate, 50 μM 2-mercaptoethanol (Gibco), 100 IU·mL^-1^ of penicillin, 100 μg·mL^-1^ of streptomycin (Bio-Concept) and 2 mM L-glutamine solution (Bio-Concept)) supplemented with 3000 IU·mL^-1^ IL-2 (Proleukin) which was gradually depleted over 48 hours to end up at 100 IU·mL^-1^. The day before the assay, 5-10×10^6^ tumor cells were plated for overnight adhesion in a 24-wells plate. The next morning, TILs were cocultured with autologous tumor cells at a ratio of 1:1 for 6 hours and 0.5–1.0×10^6^ TILs were kept unstimulated as a background control. Cells were recovered and the upregulations of CD137 was evaluated by staining with anti-41BB/CD137 (Miltenyi), anti-CD4 (BD Biosciences), anti-CD8 (Biolegend) and Far Red Dead Cell Stain (Invitrogen) for 20-30 min at 4°C. After washing, samples were finally diluted in 500 µL of PBS and the tumor-reactive fraction sorting was performed using either a Sony SH800 Cell Sorter (Sony) or a BD FACS Melody (BD Biosciences). Tumor-reactive CD4^+^/CD8^+^ TILs were FACS sorted based on CD137 upregulation and purified cells were used for bulk TCR-sequencing. The non-tumor-reactive fraction (i.e. CD137-negative) was also sorted and TCR-sequenced as control. Data were then analyzed using FlowJo software (LLC) and background (i.e. TILs alone) was subtracted for downstream analyses.

### Bulk TCR α and β sequencing

mRNA was isolated using the Dynabeads mRNA DIRECT purification kit (Life Technologies) and was amplified using the MessageAmp II aRNA Amplification Kit (Ambion) with the following modifications: *in vitro* transcription (IVT) was performed at 37°C for 16 hours. First, strand cDNA was synthesized using the Superscript III (Thermo Fisher Scientific) and a collection of *TRAV/TRBV*-specific primers. TCRs were then amplified by PCR (20 cycles with the Phusion from New England Biolabs, NEB) with a single primer pair binding to the constant region and the adapter linked to the *TRAV/TRBV* primers added during the reverse transcription. A second round of PCR cycle (25 cycles with the Phusion from NEB) was performed to add the Illumina adapters containing the different indexes. The TCR products were purified with AMPure XP beads (Beckman Coulter), quantified and loaded on the MiniSeq instrument (Illumina) for deep sequencing of the TCRα/TCRβ chain. The TCR sequences were further processed using *ad hoc* Perl scripts to (i) pool all TCR sequences coding for the same protein sequence; (ii) filter out all out-frame sequences; and (iii) determine the abundance of each distinct TCR sequence. TCR sequences with a single read were not considered for analysis. This sensitive in-house bulk TCR-Seq method is further described in Genolet *et al.*^61^. To normalize and compare TCR repertoire metrics, blood and expanded TIL repertoires were performed on 1’000’000 cells, while 0.5 µg of RNA was used from tumor samples.

T-cell subtypes (CD4^+^, CD8^+^ or undetermined) annotation of TCR sequences found in bulk TCR-seq was performed as follows: every TCRβ chain was mapped to the annotated and integrated scRNA-seq/scTCR-seq data using all time points (T0, ACT product and T30) (described below). In frequent cases where a TCRβ chain was mapped to different T-cell subtypes, the preponderant phenotype was attributed to the chain. In case of *ex aequo*, the TCR sequence was left as “undetermined”.

### Selection of tumor-reactive and clonally expanded TCRs

For the tumor-reactive repertoire sorted from the ACT product, the top 1-6 clonotypes were selected from the CD137^+^-sorted TCRβ sequencing for each patient and the corresponding alpha was found based on scTCR-seq (**Supplementary Table 2**). Potential tumor-reactive clonotypes had to satisfy the following rules: the percentage of the TCRβ chain in the CD137-positive fraction must be 1.5 time bigger than the cognate percentage in the CD137-negative fraction and bigger than its percentage in bulk ACT product repertoire. For baseline tumors, the top 10 most expanded TCRs were extracted from scTCR-seq before processing and cloned for tumor reactivity screening. These TCRs were then mapped on processed data and TCRs were ranked according to their frequency. For ACT products, the top 1-3 most expanded TCRs were extracted from scTCR-seq and cloned for tumor reactivity screening. The tumor reactivity screening results of additional TCRs identified through NeoScreen^62^ or through a screening for viral specificities (using the CEF pool, JPT) are also depicted in **Supplementary Table 2** and used for further analysis.

### Design of DNA constructs and *in vitro* transcription of RNA for TCR cloning

DNA sequences coding for full-length TCR chains were codon optimized and synthesized by GeneArt (Thermo Fisher Scientific) or Telesis Bio as strings. Each DNA sequence included a T7 promoter upstream of the ATG codon while human constant regions of α and β chains were replaced by corresponding homologous murine constant regions. Strings served as templates for the IVT and polyadenylation of RNA molecules as per manufacturer’s instructions (HIScribe T7 ARCA mRNA kit, NEB). Polyadenylation and integrity were assessed by gel electrophoresis in denaturing conditions and RNA was quantified with a Qbit fluorometer (Thermo Fisher Scientific). Purified RNA was resuspended in water at 1-10 μg·mL^-1^ and stored at -80°C until used.

### TCR cloning and tumor reactivity validation

To interrogate antitumor reactivity, TCRαβ pairs were cloned into recipient activated T cells or Jurkat cell line (TCR/CD3 Jurkat-luc cells (NFAT), Promega, stably transduced with human CD8alpha/beta and TCRalpha/beta CRISPR-KO). Autologous or HLA-matched allogeneic PBMCs were plated at 10^6^ cells·mL^-1^ in 48-well plates in R8 medium supplemented with 50 IU·mL^-1^ IL-2 (Proleukin). T cells were activated with Dynabeads Human T Activator CD3/CD28 beads (Thermo Fisher Scientific) at a ratio of 0.75 beads: 1 total PBMC. After 3 days of incubation at 37°C and 5% CO_2_, beads were removed and activated T cells rested for two extra days before electroporation or freezing. For the transfection of TCRαβ pairs into T cells and Jurkat cells, the Neon electroporation system (Thermo Fisher Scientific) was used.

Briefly, cells were resuspended at 15-20×10^6^ cells·mL^-1^ in buffer R (buffer from the Neon kit), mixed with 300-500 ng of TCRα chain RNA together with 300-500 ng of TCRβ chain RNA and electroporated with the following parameters: 1600V, 10ms, 3 pulses and 1325V, 10ms, 3 pulses, for activated PBMCs and Jurkat cells respectively. Electroporated cells were either incubated for 5-6 hours at 37°C before being used in co-culture assays. To assess antitumor reactivity of TCRs, 10^5^ TCR RNA-electroporated T cells and 2×10^4^ to 10^5^ autologous tumor cells pre-treated or not with IFNγ were cocultured in 96-wells plates. After overnight incubation, cells were recovered and the upregulation of CD137 in T cells was evaluated by staining with anti-41BB/CD137 (Miltenyi), anti-CD3 (Biolegend or BD Biosciences), anti-CD4 (BD Biosciences), anti-CD8 (BD Biosciences) and anti-mouse TCRβ-constant (Thermo Fisher Scientific) and with Aqua viability dye (Thermo Fisher Scientific). For the luciferase assay, 5×10^4^ Jurkat cells were cocultured with 1×10^4^ to 5×10^4^ autologous tumor cells pre-treated or not with IFNγ in 96-well plates. After overnight incubation, the assay was performed using the Bio-Glo Luciferase Assay System (Promega). The following experimental controls of TCR transfection were included: MOCK (transfection with water), a control TCR (irrelevant crossmatch of a TCRα and β chain). FACS samples were acquired with LSRFortessaTM (BD Bioscience) or IntelliCyt iQue® Screener PLUS (Sartorius) flow cytometers and FACS data analyzed with FlowJo v10 (TreeStar) and ForeCyt v6.2.6752 (Intellicyt). Luminescense was measured with a Spark Multimode Microplate Reader (Tecan). Validation of non-tumor-reactivity of TCRαβ pairs required the percentage of CD137 expression or luminescent units after tumor challenge to be equal or lower than background level in all control conditions and the validation of the pairing by scTCR-seq data.

### Bulk RNA extraction and sequencing

Total RNA was extracted from available snap frozen tissues (T0 and T30). Samples were lyzed in TRIzol reagent (Invitrogen) using a tissue lyzer (Qiagen) and the RNA was purified with the RNeasy kit (Qiagen). RNA quality was assessed with a Fragment Analyzer (Agilent) and Nanodrop spectrophotometer (Thermofisher). Quantification was performed with the Qubit HS RNA assay kit (Invitrogen). 0.5 μg of RNA was kept for bulk TCR-sequencing (see corresponding method). RNA sequencing libraries were prepared using the Illumina TruSeq Stranded Total RNA reagents according to the protocol supplied by the manufacturer and sequenced using HiSeq 4000. Illumina paired-end sequencing reads were aligned to the human reference GRCh37.75 genome using *STAR* aligner (version 2.6.0c) and the 2-pass method as briefly follows: the reads were aligned in a first round using the *--runMode alignReads* parameter, then a sample-specific splicejunction index was created using the *--runMode genomeGenerate* parameter. Finally, the reads were aligned using this newly created index as a reference. The number of counts was summarized at the gene level using *htseq-count* (version 0.9.1). The Ensembl ID were converted into gene symbols using the *biomaRt* package and only protein-coding, immunoglobulin and TCR genes were conserved for the analysis. Read counts were normalized into reads per kilobase per million (RPKM) and log_2_ transformed after addition of a pseudo-count value of 1 using the *edgeR* R package. As the data was processed in three different batches, we applied a batch correction algorithm using the *ComBat* function of the *sva* R package by using the patient origin as a covariate in the model.

### FACS sorting, encapsulation, and single-cell library construction

The samples of the infused T cell product were thawed, and TILs undergone overnight recovery in R8 medium supplemented with 3000 IU·mL^-1^ IL-2 (Proleukin). TILs were resuspended in PBS + 0.04% BSA and DAPI (Invitrogen) staining was performed. Live cells were sorted with a BD FACS Melody sorter and manually counted with a hemocytometer and viability was assessed using Trypan blue exclusion. For baseline tumor samples and post-ACT biopsies, either frozen pieces were thawed in RPMI + 20% FBS on the day of the assay or fresh tissue was directly chopped upon reception. Tissue was dissociated in RPMI + 2% Gelatin (Sigma-Aldrich) + 200 IU·mL^-1^ Collagenase I (ThermoFisher Scientific) + 400 IU·mL^-1^ Collagenase IV (ThermoFisher Scientific) + 5 IU·mL^-1^ Deoxyribonuclease I (Sigma-Aldrich) + 0.1% RNasin Plus RNase Inhibitor (Promega) for 15-30 min at 37°C. Digested cells were filtered using a 70 μm strainer and resuspended in PBS + 1% Gelatin + 0.1% RNasin. Cells were manually counted then stained with 50 µM·mL^-1^ of Calcein AM (Thermo Fisher Scientific) and FcR blocked (Miltenyi Biotec) for 15 min at RT. Surface staining with anti-CD45 (BioLegend) was performed for 20 min at 4°C. Cells were resuspended in PBS + 0.04% BSA (Sigma-Aldrich) and DAPI (Invitrogen) staining was performed. For post-infusion biopsies, due to small cell numbers, after counting, cells were directly stained with anti-CD45 (BioLegend) and viability stained with PI and Hoechst. For baseline tumors, 40’000 CD45^+^ cells were sorted on a MoFlo AstriosEQ (Beckman Coulter) and collected in 0.2mL PCR tubes containing 10 μL of PBS + 0.04% BSA + 0.1% RNasin. For post-infusion biopsies, depending on the percentage of CD45^+^ fraction in the sample, two sorting strategies were adopted. If the CD45^+^ fraction was inferior to 75% of total viable cells, 40’000 CD45^+^ and 40’000 CD45^-^ viable cells were collected in two separated tubes. If the CD45^+^ fraction was superior to 75%, 40’000 total viable cells were sorted. After sorting, cells were manually counted with a hemocytometer and viability was assessed using Trypan blue exclusion. In cases when CD45^+^ and CD45^-^ were sorted separately, cells were mixed with a ratio of 70% CD45^+^ and 30% CD45^-^. TILs from ACT products and *ex vivo* CD45^+^ cells from tumors were resuspended at a density of 600-1200 cells µL^-1^ with a viability of >90% and subjected to a 10X Chromium instrument for single-cell analysis. The standard protocol of 10X Genomics was followed and the reagents for the Chromium Single Cell 5’ Library and V(D)J library (v1.0 or v1.1 Chemistry) were used. 15’000 cells were loaded per sample when possible (or the total number of sorted cells if < 15’000 cells). Using a microfluidic technology, single-cell were captured and lysed, mRNA was reverse transcribed to barcoded cDNA using the provided reagents (10X Genomics). 14 PCR cycles were used to amplify cDNA and the final material was divided into two fractions: first fraction was target-enriched for TCRs and V(D)J library was obtained according to manufacturer protocol (10X Genomics). The second fraction was processed for 5’ gene expression library following the manufacturer’s instruction (10X Genomics). Complementary DNA and library quality were examined on a Fragment Analyzer (Agilent) and quantification was performed with the Qubit HS dsDNA assay kit (Invitrogen). Barcoded V(D)J libraries and GEX libraries were pooled and sequenced by an Illumina HiSeq 4000 sequencer.

### Data processing of scRNA-seq and scTCR-seq libraries

Tumors (baseline, T0 and post-ACT biopsy, T30) and ACT product (ACTP) samples were analyzed separately. T0 samples involved 15 samples representing 13 patients (two patients had two different tumor site biopsies), T30 samples involved 7 samples from 7 patients and 13 ACT product samples were analyzed (one per patient). T0 samples involving two different sites for the same tumor were pooled for subsequent analyses unless otherwise mentioned. The scRNA-seq reads were aligned to the GRCh38 reference genome and quantified using *cellranger* count (10X Genomics, version 4.0.0). Filtered gene-barcode matrices that contained only barcodes with unique molecular identifier (UMI) counts that passed the threshold for cell detection were used and processed using the *Seurat* R package version 4.0.1. For tumor samples (T0 and T30), the number of genes expressed per cell averaged 1’458 (median: 1’240) and the number of unique transcripts per cell averaged 4’206 (median: 2’926). Low quality cells exhibiting more than 10% of mitochondrial reads were discarded from the analysis. After removing the low-quality cells, we obtained 93’522 cells altogether (61’991 for T0 and 31’531 for T30). For ACT product sample analysis, the number of genes expressed per cell averaged 1’883 (median: 1’775) and the number of unique transcripts per cell averaged 4’899 (median: 4’208). We obtained 69’923 cells altogether (5’379 (mean) cells +/- 1’486 (s.d.) per sample) and were left with 65’757 cells after removal of low-quality cells (exhibiting more than 10% of mitochondrial reads). scTCR-seq (VDJ) data were aligned to the same human genome using the *cellranger vdj* (10X Genomics, version 3.1.0). Only true cells (with a “True” label in the “is_cell” column of the all_contig_annotations.csv file) were kept for further analyses. Cells from the VDJ sequencing were mapped to the scRNA-seq data (GEX). We found that 65.4% of the T cells from the tumor had a mapped TCR β-chain (58.4% for TCR α-chain). For the ACT product, 90.5% of the T cells had a mapped TCR β-chain (85.5% for TCR α-chain).

### Single-cell clustering analysis

The data was processed using the *Seurat* R package (version 4.0.1) as follows briefly: counts were log-normalized using the *NormalizeData* function (scale.factor=10’000) and then scaled using the *ScaleData* function by regressing the mitochondrial and ribosomal rate of read contents, the number of read count per cell (nCount_RNA, only for T0 and T30 tumor data), and cell cycle parameters represented by S phase and G2/M phase scores (computed using the *CellCycleScoring* function with the list of genes provided internally in the *Seurat* package). Dimensionality reduction was performed using the standard Seurat workflow by principal component analysis (*RunPCA* function) followed by tSNE and UMAP projection (using the first 75 principal components, PCs). The k-nearest neighbors of each cell were found using the *FindNeighbors* function run on the first 75 PCs, and followed by clustering at several resolutions ranging from 0.1 to 10 using the *FindClusters* function.

### Cell type annotation

#### Main cell-lineage annotation

The annotation of main cell types was performed using the combination of several different methods: i) differential gene expression of clusters at different resolutions using the Seurat *FindAllMarkers* function (with the default Wilcoxon statistical testing parameter) followed by literature curation; ii) investigation of the expression of known canonical gene markers for melanoma cells (*MLANA, PRAME, SOX10, S100B*), immune cells (*PTPRC*), T cells (*CD3E, CD8A, CD8B, CD4*), B cells (*CD79A, MS4A1*), CAFs (*DCN, FAP*), endothelial cells (*PECAM1, VWF*), plasma cells (immunoglobulins), Myeloid cells (*CD68, HLA-DRA, LYZ, CD86*); iii) automated annotation using the *singleR* package and using gene expression centroids (average per cell described cell populations) derived from several studies: Yost *et al.*^34^, Guo *et al.*^63^, Zhang *et al.*^64^, Oliveira *et al.*^21^, Zilionis *et al.*^65^ and the human primary cell atlas (HPCA) reference as inferred in the *singleR* package; iv) the presence or absence of mapping scTCR-seq data indicating T cells versus non-T cells. Once major clusters were annotated into broad cell types (T cells, B cells, malignant cells, myeloid cells, CAFs and endothelial cells) further intra-cluster investigations allowed finer cellular subtypes annotations (detailed version is disclosed in Barras *et al.*^25^).

#### Main cell subtype annotation of T cells and NK cells in tumor and ACT product

For finer annotation of the T/NK cells, the cells were first classified as CD8^+^, CD4^+^, double-negative (DN; CD8^-^/CD4^-^), double-positive (DP; CD8^+^/CD4^+^ doublets), NK cells and Tγδ as follows: cells with non-null expression of *CD8A* and null expression of *CD4* were defined as CD8^+^ (and *vice-versa* for CD4^+^). Cells showing non-null expression of both genes were first classified as DP, then as doublets of CD4^+^ and CD8^+^ T cells as the average number of genes expressed per cell equaled close to the double of CD4^+^ or CD8^+^ T cell singlets. Due to notorious dropout events in single-cell data, cells lacking the expression of both markers were classified as follows: if a cell belongs to a cluster (considering a high resolution of 10) in which the 75^th^ percentile expression of *CD8A* was higher than its 75^th^ percentile expression of *CD4*, it was classified as CD8^+^ (and *vice-versa* for CD4^+^). If the 75^th^ percentile expressions of both markers equal 0, the cells were classified as DN. Finally, cells with an average expression scores of all *TRG* and *TRD*-related genes higher than 0.5 (cutoff established after histogram visual inspection) were assigned to be Tγδ cells. We also characterized the T-cell repertoire of CD45^+^-sorted cells by scTCR-seq and compiled additional Tγδ cells for which TCR gamma or delta chains were found. In the general clustering of all CD45^+^-sorted cells, NK cells were clustering close to the CD8^+^ T cells and were annotated as NK1 and NK2 using the Zilionis *et al*. (*26*) centroid annotation by the *singleR* function from the *singleR* package.

#### High-resolution CD4^+^ and CD8^+^ T cell subtypes in the tumors

The clustering of CD4^+^ T cells was obviously formed by three distinct clusters whose gene markers were indicating CD4 CXCL13 (T follicular-helper) cells (*CXCL13^+^*, *CD40LG^+^*, *BCL6^+^*, *CD200^+^*), CD4 Tregs (*FOXP3^+^*, *CTLA4^+^*, *IL2RA^+^*) and CD4 T helper 1 (Th1) cells (*IL7R^+^*, *SELL^+^*, *LEF1^+^*).

For the CD8^+^ T cell subtyping, CD8^+^ T cells and NK cells were integrated by sample and by removing the TCR genes in order to prevent clustering based on clonotypes (due to TCR gene expression) as explained in Barras *et al.*^25^. We used several methods to annotate the clusters:

1. We first performed differential gene expression and differential regulon/TF activity (see below) analysis which were computed using the *FindAllMarkers* function with a 0.7 resolution.
2. We computed signature score (using the *AUCell* R package) for cytotoxicity (*CCL3*, *CCL4*, *CST7*, *GZMA*, *GZMB*, *IFNG*, *NKG7*, *PRF1*), exhaustion (*CTLA4*, *HAVCR2*, *LAG3*, *PDCD1*, *TIGIT*) and naiveness/memory (*CCR7*, *LEF1*, *SELL*, *TCF7*) taken from the Table S3 of Jerby-Arnon *et al*. list of genes^66^. We extracted the gene list from Figure 7 of Andreatta *et al*.^67^ as specific markers. We used the prediction from the annotation coming from public studies, namely Yost *et al*.^34^, Guo *et al*.^63^, Zhang *et al*.^64^ and Oliveira *et al*.^21^. The annotation of high-resolution clusters was performed as follows:

1. A cluster displaying elevated levels of *CCR7*, *LTB*, *SELL* and *IL7R* gene expression, high TCF7 and LEF1 regulon activity, high concordance with Oliveira *et al*.^21^ predictions of naive cells was then named CD8 naive-like.
2. Two clusters were displaying signs of stress or apoptosis: one with elevated levels of mitochondrial-related genes (gene names starting by *MT-*), expression of the *MALAT1* gene and lower number of genes expressed per cell was then named CD8 low-quality and another one with overexpression of heat-shock protein (HSP) genes and *DNAJ*-related genes which we named CD8 HSP.
3. A cluster displaying concomitant expression of *CD8A* and *FOXP3* genes, without no signs of higher number of genes per cell (which could have otherwise highlighted doublets of CD8^+^ T cells and CD4 Tregs) and high concordance with Oliveira *et al*.^21^ predictions of Treg-like cells was then named CD8 FOXP3.
4. A cluster was obviously driven by the high expression of type-I interferon genes (*ISG15*, *MX1*, *IFI16*, *IFIT3*, *IFIT1*, *ISG20*, *OAS1*) and was then named CD8 ISG.
5. A small cluster was found very close to the one of NK cells and displaying *CD8A* expression with co-expression of NK cells marker such as *KLRC2* and *KLRD1*, and high concordance with Oliveira *et al*.^21^ predictions of NK-like cells was then named CD8 NK-like.
6. Another small cluster that was captured at a finer resolution of 2, that displayed high expression of *CX3CR1* and *FGFBP2*, high levels of cytotoxic and low levels of exhaustion signatures, high concordance with Guo *et al*.^63^ and Zhang *et al*.^64^ predictions of CD8_CX3CR1 cells was then named CD8 CX3CR1.
7. Four clusters with similar gene expression profiles, displaying high expression of *GZMK* (a marker for effector memory cells according to Andreatta *et al*.^67^, modest cytotoxic and low exhaustion signature levels, high concordance with Oliveira *et al*.^21^ predictions of CD8 Effector-Memory (EM) cells were then combined and named CD8 EM-like.
8. Two clusters showing obvious signs of exhaustion with high levels of *HAVCR2* and *PDCD1* expression, high exhaustion signature levels, and elevated EOMES regulon activity were first classified as exhausted T cells then subclassified as follows: one cluster has higher *HAVCR2*, a typical sign of late exhaustion, and high concordance with Oliveira *et al*.^21^ predictions of Terminal-Exhausted cells and was then named CD8 Tex; and the other cluster displayed higher levels of *TOX*, *XCL1*, *XCL2* and *CRTAM*, which are markers of precursor exhaustion according to Andreatta *et al*.^67^, lower activity of TBX21 regulon and high concordance with Oliveira *et al*.^21^ predictions of Progenitor-Exhausted cells and was thus named CD8 Pex.
9. Finally, three clusters were driven by proliferation markers. Since proliferation is not a T-cell state in itself, we generated an average expression profile (centroid) for our CD8^+^ T cell subsets and predicted the T-cell state of proliferating CD8^+^ T cells by automated annotation using this centroid and the *singleR* R package. The full list of differentially expressed genes and TFs per cell types appears in Barras *et al.*^25^ and was computed using the *FindAllMarkers* function from the *Seurat* package.

#### Cell annotation of the ACT product

As explained above, the ACT product was annotated for the main cell types (T cells, NK cells and myeloid cells) and main T cell subtypes (CD8-positive, CD4-positive, DN, DP and Tγδ populations). CD4 Tregs in the ACT product were annotated by tracking TCR sequences from baseline tumors corresponding to CD4 Tregs and annotating ACTP cells with the same TCR sequences as Tregs.

Unique ACT product TCR sequence often matches to several tumor *in situ* cells belonging to different cell states. Thus, in order to assess the contribution of intratumoral CD8^+^ T cell subtypes to ACT products and to take into account the initial T0 cell state frequency in a given patient, we first overlapped the scTCR-seq from ACTP with the scTCR-seq repertoire of the T0 tumor (defining TCR sequence as the concatenation of the two TCRβ chains). ACT product cells without any matching TCR sequence in T0 were labeled as “not_in_T0”. The TCR sequences of the other cells were reverse tracked in the T0 tumor. For each TCR sequence, the number of corresponding T0 cells per cell state was normalized by the overall frequency of these cell states in a given patient. The cell state with the highest value was attributed to this TCR sequence in the ACT product.

Finally, analysis of *pure* CD8^+^ T cell TCRs’ (TCR sequence for which only one cell state was found in the matched T0 scTCR-seq data) expansion and transcriptome was allowed by defining *pure* clonotypes as clones composed only by cells in the same *in situ* CD8^+^ T cell state.

### Overlap, diversity and clonality analyses of TCR repertoires

Bulk TCR-seq data were used to compute richness and clonality metrics previously described^35^. Only TCRβ chains were considered for the computations. Richness was assessed by the number of unique TCR sequences present in the repertoire. The clonality was described by the Shannon Entropy or 1-Pielou’s evenness, according to the formulas described in Chiffelle *et al*^35^. The overlap of TCR repertoires was computed as the cumulative frequencies of the shared sequences between two repertoires within each repertoire.

### Gene signature analysis

For single-cell data analysis and unless otherwise mentioned, gene signature scores were computed using the *AUCell* package. For bulk RNA sequencing data and bulked scRNA-seq data (i.e. pseudo bulk analysis of ACTP), gene signature scores were computed using single-sample geneset enrichment analysis (ssGSEA) method as inferred in the *gsva* function from the *GSVA* R package. Individual gene signatures were taken from the source indicated in the figure, we used the cytotoxicity and exhaustion signatures from Jerby-Arnon *et al.*^66^ under the names “CYTOTOXIC T CELL (SPECIFIC MARKERS)” and “EXHAUSTED T CELL (SPECIFIC MARKERS)”, respectively. We also used the Reactome collections taken from the MSigDB portal (http://www.broadinstitute.org/gsea/msigdb) by isolating the pathway names starting with the “REACTOME_” term in the C2 collection. Gene signatures used in the analysis of **Figure 5B** were identical to those used in Barras *et al*.^25^.

When tracking gene signature scores from T0 to ACT product, then to T30, we used bulk RNA-seq data for T0 and T30 and we average-pseudobulked ACT product scRNA-seq data in order to allow comparing all time points and to maximize the number of patients analyzed (since T30 bulk RNA-seq data was available for 10 patients while scRNA-seq data was available only for 7 patients out of 13).

Gene signatures specific for cell subsets identified in our single-cell CD45^+^ cell-sorted data were derived by performing differential gene expression using the *FindAllMarkers* function from the *Seurat* package by using the default parameters. The genes kept as specific to a cell subset were the 20 most non-TCR-related differentially expressed (lowest *p*-values) genes.

### Signature score and prediction of tumor reactivity

The CD8^+^ T cell tumor reactivity signature was obtained by performing differential gene expression between tumor-reactive (*n*=1499 cells) and non-tumor-reactive (*n*=1354 cells) CD8^+^ T cells (**Supplementary Table 2**) using linear regressions as inferred in the *lmFit* function of the *limma* R package. TCR-related genes and genes with average expression values lower than 0.3 were discarded from the differentially expressed gene list. After differential expression analysis between defined tumor-reactive and non-reactive CD8^+^ T cells, the tumor reactivity signature was obtained by selecting the 20 most significantly non-TCR-related upregulated genes (**Supplementary Table 3**). We computed the statistical enrichment of external published directed (upregulated and downregulated genes) signatures from Oliveira *et al.*^21^, Lowery *et al.*^16^, He *et al.*^17^ and Zheng *et al.*^18^ in our tumor-reactive gene signature by using the *hypeR* function from the *hypeR* R package (default parameters) as given by the false-discovery rate. Finally, the tumor-reactive score for every cell was obtained by using the *AUCell* signature score method. An *in vitro* validation of this signature score was carried on in two patients with a low number of tumor-reactive clones among the top 10 expanded from the tumor (patients #4 and #5). 5-6 TCRs with high and low scores were tested per patient resulting in a discriminative probability of AUC = 0.876, p = 0.0024. In order to interpret the continuous tumor-reactive scores, TCR clones were stratified into discrete tumor reactivity categories. As a starting point, scTCR-seq data of CD8^+^ T cells were grouped by TCR clones (both α and β chains) by averaging the tumor reactivity scores. The clonotypes were subsequently divided into equally distributed categories with increasing tumor-reactive scores conveniently named *Low*, *Medium* and *High*. Each category was equally distributed and had increasing tumor-reactive scores. This step resulted into a reference table defining the tumor reactivity status for every TCR clone that could be mapped onto the scTCR-seq and bulk-TCR-seq data. Clones with a *High* and *Low* tumor-reactive score were labeled as tumor-reactive and non-tumor-reactive, respectively. For the mapping onto single-cell data, both α and β chains were taken into consideration into a simple mapping. As for the bulk data, mapping was allowed by agglomerating the TCR clones by β chains only. To do so, we had to correct for the few cases where β chains that are expressed in combination with different α chains and were defined with different tumor reactivity status. To solve this, we interpreted the tumor reactivity as a dominant trait, thus implying the *High* category prevails the *Medium* category which prevails the *Low* category. Finally, for analysis performed with combination of scTCR-seq and bulk TCR-seq data, only the β chains were considered.

### Transcription factor activity in single-cell data

The transcription factor activity was estimated using the regulon signature of each transcription factor. Regulons were inferred using the *SCENIC* pipeline (https://scenic.aertslab.org) which integrates three algorithms (*grnBoost2*, *RcisTarget* and *AUCell*) corresponding to three consecutive steps:

Step 1: First, a gene regulatory network (GRN) was inferred from all tumors and ACT products transcriptomic together using grnBoost2, a faster implementation of the original Genie3 algorithm. grnBoost2 takes as the input scRNA-seq transcriptomics data to infer causality from the expression levels of the transcription factors to the targets based on co-expression patterns. Basically, the prediction of the regulatory network between n given genes is split into n different regression problems and expression of a given target gene was predicted from the expression patterns of all the transcription factors using tree-based ensemble methods, Random Forests or Extra-Trees. The ranking of the relative importance of each transcription factor in the prediction of the target gene expression pattern is taken as an indication of a putative regulatory event. The aggregation targets into raw putative regulons was done using the runSCENIC_1_coexNetwork2modules function from the *SCENIC* R package with default parameters.

Step 2: Co-expression modules (raw putative regulons, i.e. sets of genes regulated by the same transcription factor) derived from the GRN generated in Step 1 are refined by removing indirect targets by motif discovery analysis using cisTarget algorithm and a cis-regulatory motif database. In particular, we used hg19-500bp-upstream-7species.mc9nr.feather and hg19-tss-centered-10kb-7species.mc9nr.feather databases. The motif database includes a score for each pair motif-gene, which allows the generation of a motif-gene ranking. A motif enrichment score is then calculated for the list of transcription factor selected targets by calculating the Area Under the recovery Curve (AUC) on the motif-gene ranking using the *RcisTarget* R package (https://github.com/aertslab/RcisTarget). If a motif is enriched among the list of transcription factor targets, a regulon is derived including the target genes with a high motif-gene score.

Step 3: Finally, *AUCell* was used to quantify the regulon activity in each individual cell (https://github.com/aertslab/AUCell). *AUCell* provides an AUC score for each regulon and cell; we discarded regulons with less than five constituent elements, as the estimation of the activity of small regulons is less reliable. For the calculation of the AUC, the parameter *aucMaxRank* of the AUCell_calcAUC function was set with a fixed number of 1500 of features.

### Clone-tracked differential gene expression analyses and Reactome pathway enrichment analysis

To track differences in gene expression of clones between T0 and ACTP and between ACTP and T30, we first selected clones that were found in the three time points based on the concatenation of the two TCRβ chains (216 clones covering 5 patients) and averaged-pseudobulk gene expression per clone. We then performed clone-paired linear regressions as inferred in the *lmFit* function of the *limma* R package by using the clone identity as a covariate separately in responders and non-responders. We report the comparison of log_2_ fold-change analyses between the indicated time-points and according to response (Rs in x axis and NRs in y axis) and the significance are color-coded as indicated in the figures using an adjusted *p*-value threshold of 0.05 for significance.

After differential expression analyses, genes were then categorized according to whether they are up-, down-regulated or not regulated from T0 to ACTP then to T30, resulting in 9 possible groups. The genes in these categories were then subjected to Reactome pathway enrichment analysis as follows: these genes were first converted into Entrez ID using *mapIds* from the *AnnotationDbi* package then were subjected to Reactome enrichment analysis using the *enrichPathway* function *ReactomePA* package.

### Plotting description

Alluvial plots were generated using the *ggalluvial* R package. Heatmaps were performed using the *pheatmap* function from the *pheatmap* R package. Plotting of scRNA-seq derived UMAP was achieved using *Seurat* R package functions. Scatter plots and Fraction plots were performed using the *ggplot2* R package. All figures were reprocessed using Adobe Illustrator 2020 for esthetical purposes. Schematic figures were created with BioRender.com.

### Statistical analyses

Statistical analyses were performed using R Statistical Software and the standard *stats* library (version 4.0.1) as well as GraphPad Prism 9.1.0. The statistical tests used, and their specifications are described in the figure legends. Parametric tests, for comparing two or more groups, were applied only on normally distributed variables validated with Anderson-Darling, D’Agostino-Pearson omnibus, Shapiro-Wilk and Kolmogorov-Smirnov tests (GraphPad 9.1.0), otherwise, non-parametric tests were used.

### Data Availability

scRNA-seq, bulk RNA-seq, scTCR-seq and bulk TCR-seq data will be made publicly available in the Gene Expression Omnibus (GEO) at the time of publication under the GSE229861 accession number.

## Acknowledgements

We wish to thank all the patients, family members and staff from all the units that participated in the study.

We thank Jean-Paul Rivals and the Biobank from the Center of Experimental Therapeutics (CTE) for their assistance.

We thank the Agora Flow Cytometry Facility of the University of Lausanne and in particular Danny Labes for his help with single-cell sorting and the Lausanne Genomic Technologies Facility for scRNA-seq/scTCR-seq and RNA-seq analysis.

We also apologize to colleagues whose work could not be cited because of space limitation.

## Fundings

This study was supported by generous funding by the Ludwig Institute for Cancer Research, a grant by the state of Vaud, and a grant by Bristol Myers Squibb to GC; and grants by the Swiss National Foundation (SNF) R‘Equip (316030_205644) to DLL; and 310030_182384 and CRSII5_193749 to AH.

## Author contributions

Conceptualization: JC, AH, GC.

Methodology: JC, DB, RP, SB, MA, AA, CS, DS, AM, BM, NF, IC, FB, RG, LQ, UD, JS, JCO, DDL, AH, GC.

Investigation: JC, DB, RP, SB, MA, AA, CS, DS, AM, BM, NF, FB, RG, LQ, MBS, DDL, AH, GC.

Visualization: JC, DB, RP, SB, RG, AH.

Sample Collection and Coordination and TIL manufacturing: LK. Funding acquisition: AH, DDL, GC.

Patients treatment: AO, BNR, EG, MI, MOO, SL, KH, SZ, OM, LT. Supervision: DDL, AH, GC.

Writing – original draft: JC, AH.

Writing – review & editing: JC, DB, AH, GC.

## Competing interests

GC has received grants, research support or has been coinvestigator in clinical trials by Bristol-Myers Squibb, Tigen Pharma, Iovance, F. Hoffmann-La Roche AG, Boehringer Ingelheim. The Lausanne University Hospital (CHUV) has received honoraria for advisory services G. Coukos has provided to Genentech, AstraZeneca AG, EVIR. GC has previously received royalties from the University of Pennsylvania for CAR-T cell therapy licensed to Novartis and Tmunity Therapeutics. DDL, SB, AH, and GC are inventors on patent applications filed by the Ludwig Institute for Cancer Research Ltd on behalf also of the University of Lausanne and the CHUV pertaining to the subject matter disclosed herein and such patent applications have been licensed to Tigen Pharma SA. SZ is currently an employee of F. Hoffmann-La Roche.

OM has consulting/advisory roles for Bristol Myers Squibb, MSD, Roche, Novartis, Amgen, Pierre Fabre, Neracare; research grants from Bristol Myers Squibb, MSD, Amgen. PCL; consultant advisor or paid speaker for Bristol Myers Squibb, MSD, Novartis, Pierre Fabre, Amgen, Nektar; has received research funding from Bristol Myers Squibb, Pierre Fabre.

All the other authors have no conflict of interest to declare.

---

**Extended Data Figure 1.**
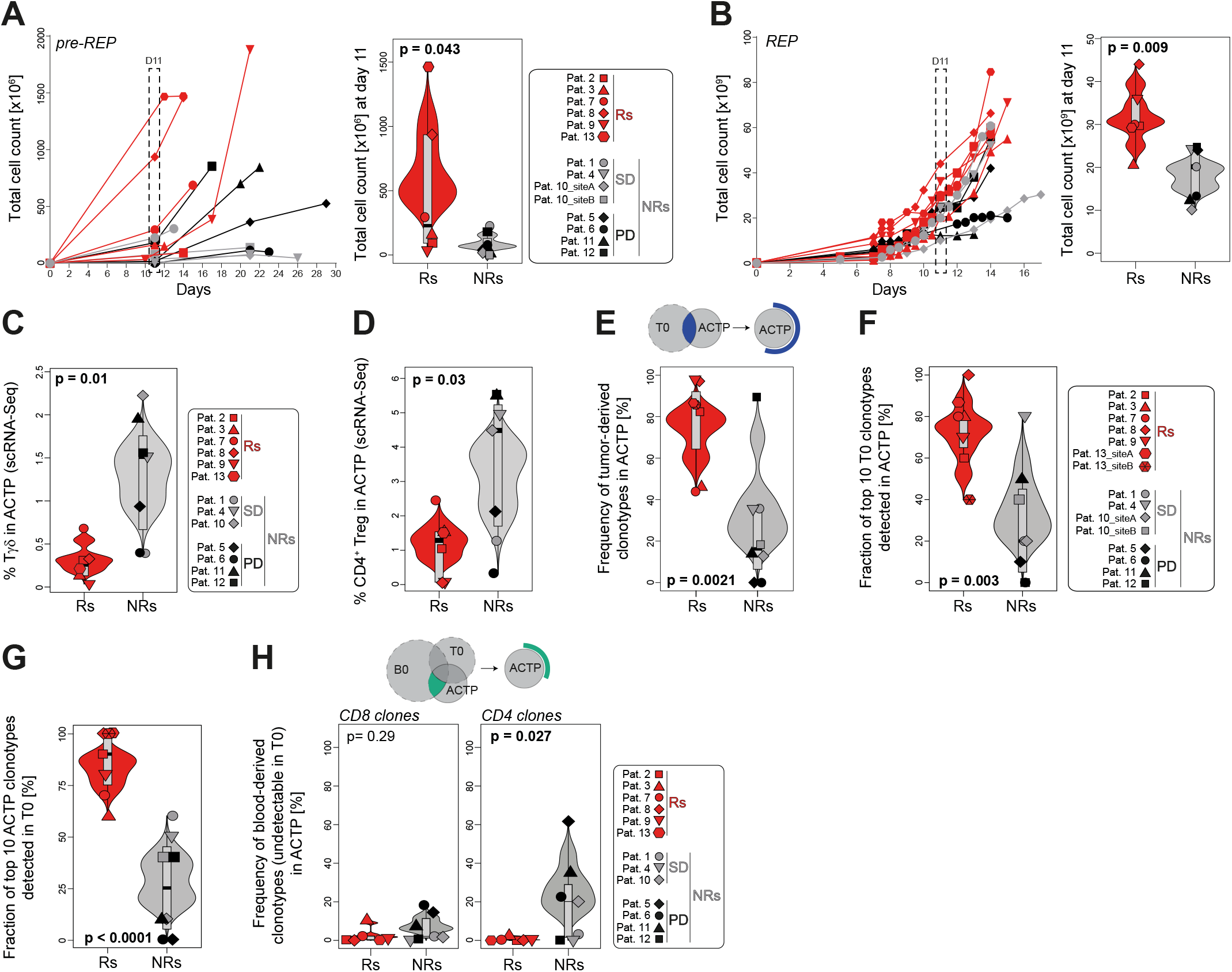
Composition of ACT products (related to Figure 1). **A-B**) Kinetics of TILs *in vitro* expansion during pre-REP (**A**) and REP (**B**) phases. Absolute cell numbers at day 11 of pre-REP (**A**) and REP (**B**) are shown on the right. Statistics were performed using two-tailed Wilcoxon and t-tests. Due to media change, patients #2, #9 and #13 were quantified at day 14, day 10 and day 12 of pre-REP, respectively, while patient #1 was quantified at day 10 of the REP. **C-D**) Percentage of Tγδ cells (**C**) and CD4^+^ Tregs (**D**) in ACTP using scRNA-seq data. CD4^+^ Tregs in ACT products (ACTPs) were identified by tracking TCR sequences of *in situ* intratumoral CD4^+^ Tregs to ACTP (see methods). Statistics were performed using two-tailed t-tests. **E**) Frequency of clones in ACTP commonly identified in tumor (T0) and ACTP. The frequency was determined using ACTP and T0 bulk TCR repertoires. Statistics were performed using a two-tailed t-test. **F-G**) Fraction of TCRs detected in cognate ACTP among the top ten dominant ones from T0 (green clones from Fig. 1J) (**F**) or detected in cognate tumors among the top ten dominant ones from ACTP (blue clones from Fig. 1J) (**G**) in Rs and NRs. Statistics were performed using two-tailed t-tests. **H**) Frequency of CD8^+^ and CD4^+^ T cell clones in ACTP identified in both blood (at baseline, B0) and ACTP but undetectable in T0. The frequency was determined using ACTP, B0 and T0 bulk TCR repertoires annotated with scRNA-seq. Statistics were performed using two-tailed Wilcoxon tests.

**Extended Data Figure 2.**
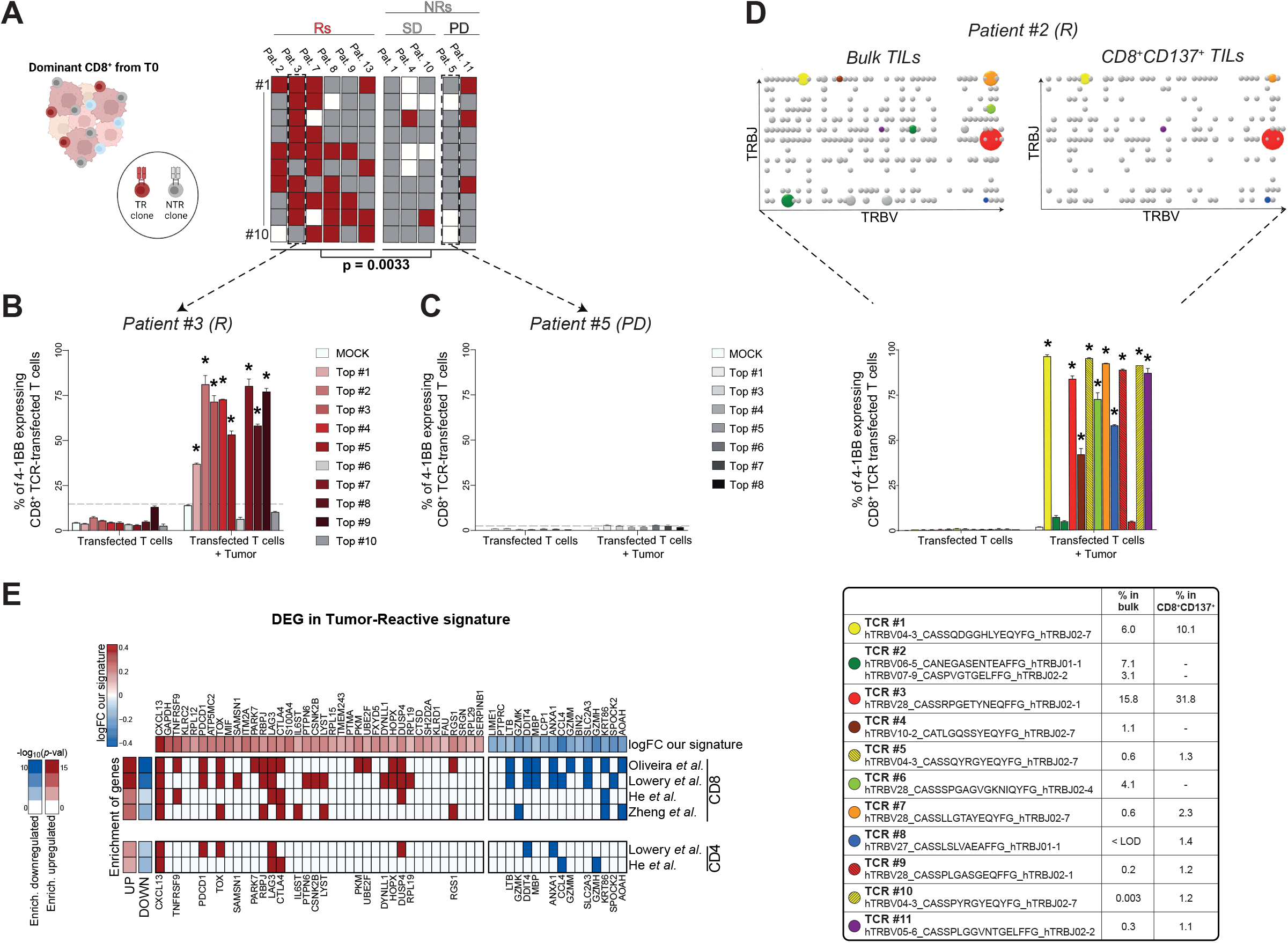
Tumor reactivity screening of clones from baseline tumors and ACT products used to build a signature of tumor reactivity (related to Figure 2). **A**) Heatmap of the distribution of validated tumor-reactive (TR) versus non-tumor-reactive (NTR) dominant CD8^+^ T cell clones from baseline tumors. Statistics were performed using a two-tailed t-test. **B-C**) Representative examples of the validation of antitumor reactivity of the dominant CD8^+^ T cell clones from baseline tumors. TCR-transfected primary CD8^+^ T cells were used and CD137 upregulation following coculture with autologous tumor cells was measured (see methods). **D**) Representative example of tumor reactivity screening of dominant clones from bulk ACTP and CD8^+^CD137^+^ tumor-reactive cells purified by FACS sorting upon coculture with autologous tumor cells for patient #2 (shown in Fig. 2A). Bulk TCR sequencing of bulk and tumor-reactive-sorted populations represented as Manhattan plots reporting TCRβ V/J recombination. Dominant TCRβ chains in bulk and tumor-reactive repertoires (see method for TCR selection) are highlighted and their frequencies are reported. The antitumor reactivity of these TCRs is validated using TCR-transfected primary CD8^+^ T cells exposed to autologous tumor cells (see methods). LOD, limit of detection. **E**) Heatmap comparing our signature of tumor reactivity with previously reported signatures found for CD4^+^ and CD8^+^ tumor-reactive T cell clones. Statistical enrichment of our signature compared to others is computed by geneset enrichment analysis as described in the methods.

**Extended Data Figure 3.**
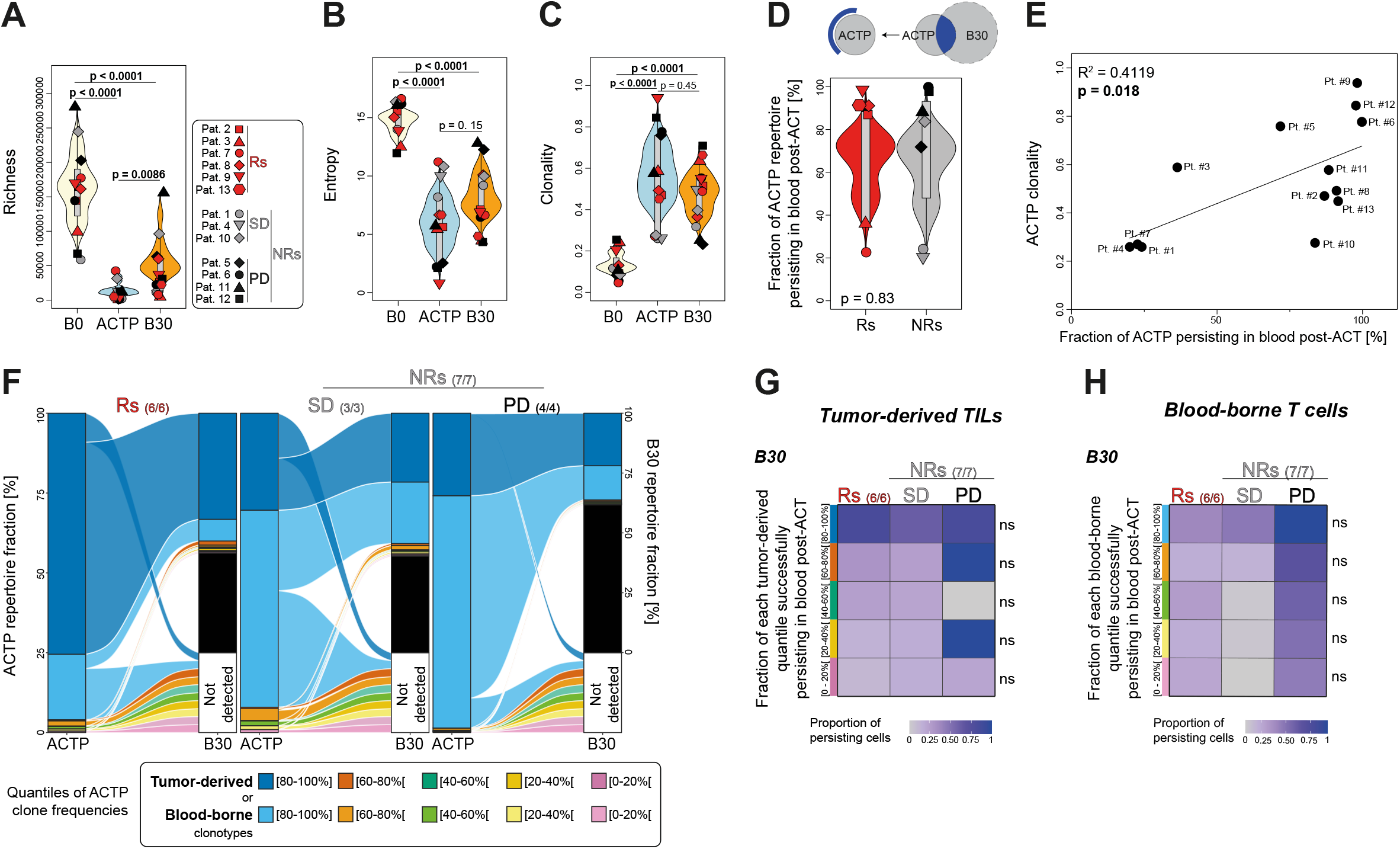
Bystander clones mostly repopulate blood circulation after ACT (related to Figure 3). **A-C**) Richness (**A**), Shannon entropy (**B**) and clonality (**C**) comparisons of bulk TCR repertoires between ACTP, pre-ACT (B0) and post-ACT (B30) blood. Statistics were performed using Wilcoxon and t-tests. **D**) Frequency of clones in ACTP commonly identified in ACTP and B30 in Rs and NRs. The frequency was determined using ACTP and B30 bulk TCR repertoires. Statistics were performed using a two-tailed Wilcoxon test. **E**) Correlation between ACTP clonality and the fraction of ACTP persisting in blood post-ACT. Analysis was performed using linear regression (R^2^ and *p*-value are shown). **F**) Fractions of ACTP repertoires (separated between tumor-derived – detected in T0 – and blood-borne – not detected in T0 but detected in B0 – clones divided into five color-coded quantiles of equivalent TCR proportions to normalize for clonality) identified in cognate post-ACT blood repertoires. ACTP clones undetectable in B30 fall in the white box (not detected). B30 clones undetectable in ACTP are represented in black. **G-H**) Heatmap showing, for each of the five quantiles of ACTP tumor-derived (**G**) or blood-borne (**H**) clones, the proportion of cells successfully persisting (i.e. identified) in B30. Statistics between Rs and NRs were performed using two-tailed t-tests.

**Extended Data Figure 4.**
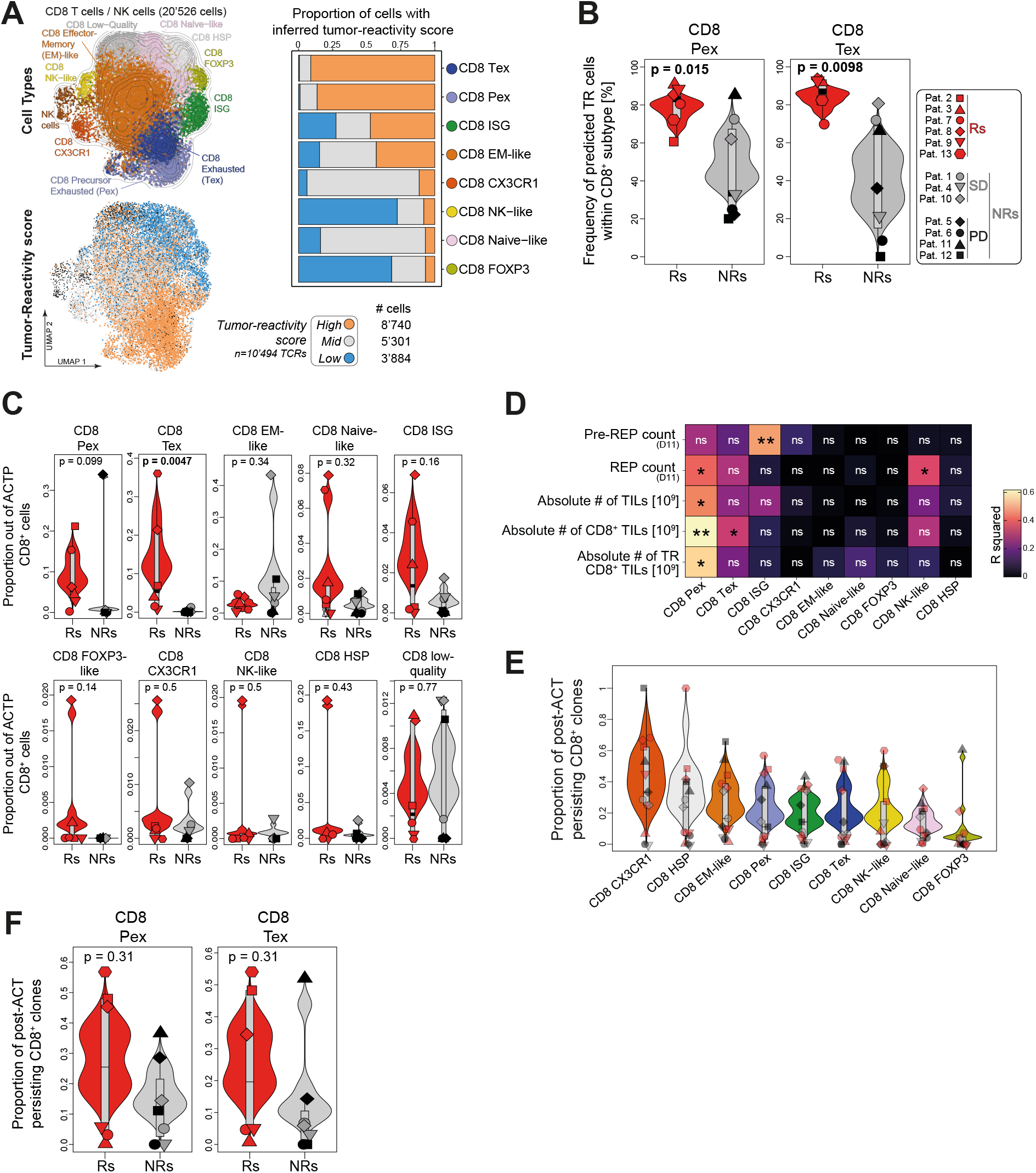
Responders’ ACTPs are mostly contributed by exhausted cells endowed with preferential infiltration capacity over blood persistence (related to Figure 4). **A**) UMAP of CD8^+^ T cells in baseline tumor scRNA-seq data color-coded according to their CD8^+^ T cell subtype (top) and to their predicted tumor reactivity status (*High*, *Mid* and *Low*) (bottom). The corresponding number of annotated cells is shown below. Proportions of cells falling in each of the predicted tumor reactivity status (*High*, *Mid* and *Low*) for the different CD8^+^ T cell subtypes identified in T0 scRNA-seq data are shown on the right. **B**) Proportion of predicted tumor-reactive (*High* status) cells in intratumoral CD8^+^ Pex and CD8^+^ Tex populations in Rs and NRs. Statistics were performed using two-tailed t-tests. **C**) Contribution of each intratumoral CD8^+^ T cell subtype within CD8^+^ T cells of ACTP scTCR repertoires in Rs and NRs. Statistics were performed using two-tailed Wilcoxon and t-tests. **D**) Linear regression coefficients (R^2^) between the different intratumoral CD8^+^ T cell subtypes and the absolute numbers of cells in pre-REP (cell count at Day11), REP (cell count at Day11), in total ACTP, total CD8^+^ T cells and tumor-reactive CD8^+^ T cells in ACTP. * p<0.05, ** p<0.01. **E**) Proportion of clones from the different intratumoral CD8^+^ T cell subtypes expanding *in vitro* (i.e. detected in bulk ACTP repertoires) and successfully persisting in blood post-ACT (i.e. detected in bulk B30 repertoires). **F**) Proportion of intratumoral CD8^+^ Pex and CD8^+^ Tex populations expanding *in vitro* (i.e. detected in bulk ACTP repertoires) and successfully persisting in blood post-ACT (i.e. detected in bulk B30 repertoires) in Rs and NRs. Statistics were performed using two-tailed t-tests.

**Extended Data Figure 5.**
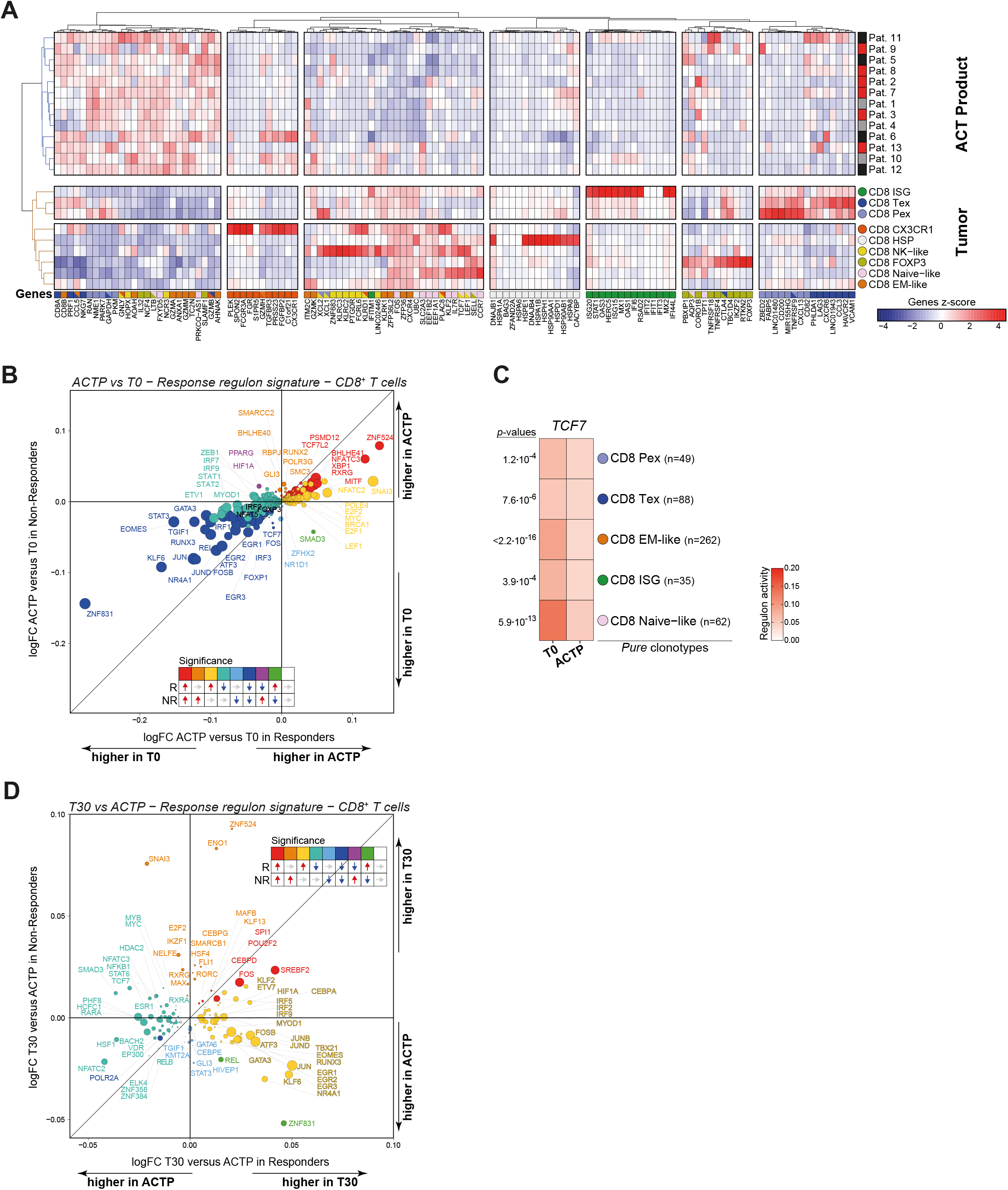
Gene and regulon changes from baseline tumors to ACT products and post-ACT tumors. (related to Figure 5). **A**) Hierarchical clustering heatmap showing the averaged-pseudobulk gene expression of canonical cell state markers of the *in situ* T0 CD8^+^ T cell subtypes and ACT products. The 15 most discriminant genes for each CD8^+^ T cell subtype are reported and color-coded. **B**) Scatter plot showing transcription factors with differential activities in CD8^+^ T cells between ACTP and T0 in Rs (x axis) and NRs (y axis). Colored dots reflect the statistical significance (adjusted *p*-value < 0.05) of each transcription factor in Rs and/or NRs and their directionality (i.e. upregulated or downregulated). **C**) Heatmap showing the averaged-pseudobulk regulon expression of *TCF7 in situ* CD8^+^ *pure* clones (i.e. containing cells found exclusively in a single state) in baseline tumors (T0) and ACT products (ACTP). The number of clones for each population is indicated in brackets. Statistics were performed using two-tailed paired t-tests. **D**) Scatter plot showing transcription factors with differential activities in CD8^+^ T cells between T30 and ACTP in Rs (x axis) and NRs (y axis). Colored dots reflect the statistical significance (adjusted *p*-value < 0.05) of each transcription factor in Rs and/or NRs and their directionality (i.e. upregulated or downregulated).

**Extended Data Figure 6.**
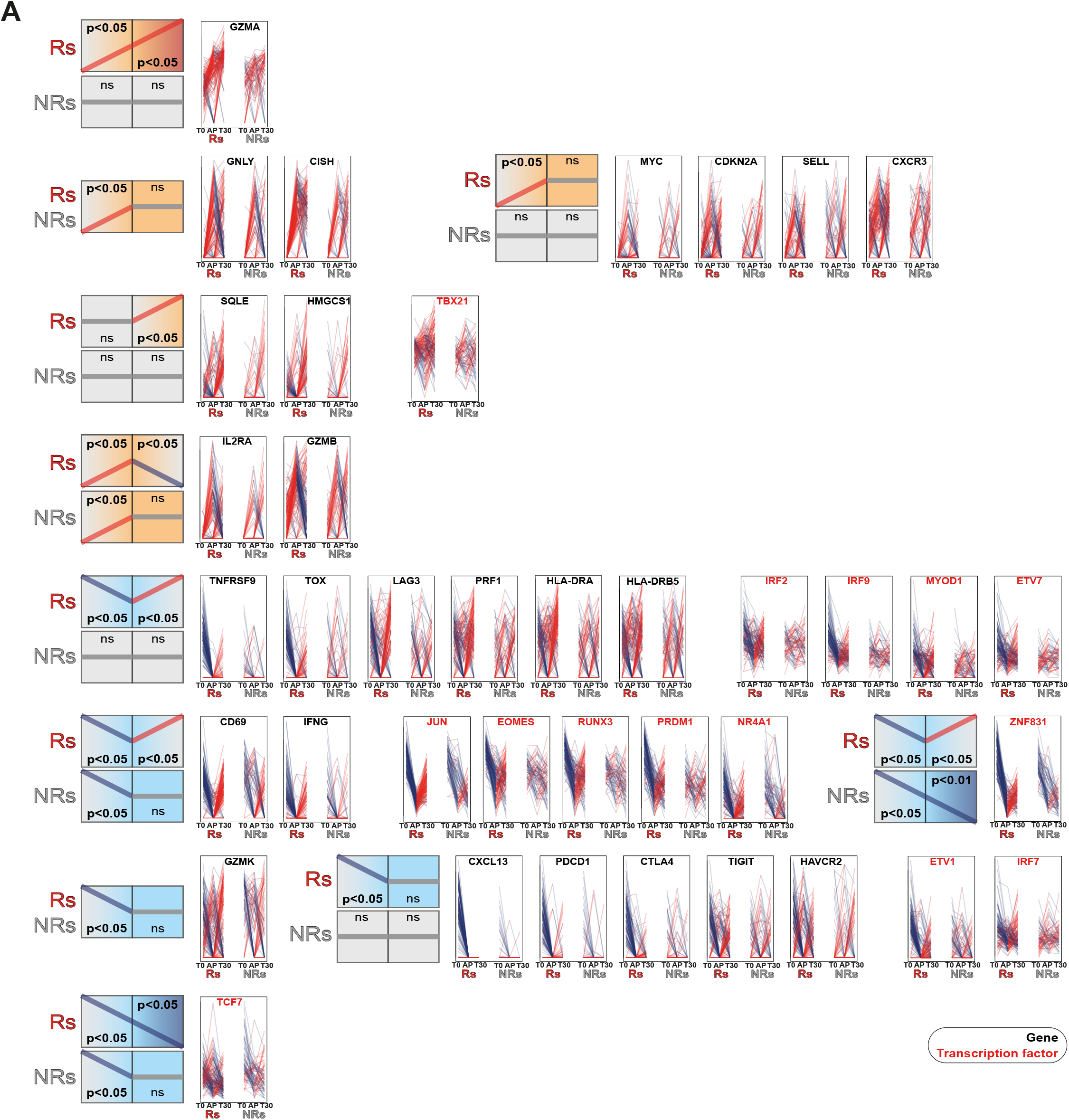
Gene and regulon changes from baseline tumors to ACT products and post-ACT tumors. (related to Figure 5). **A**) Genes and transcription factors selected from various dynamic categories in Fig. 5H. Clone averaged-pseudobulk values are shown from T0, to ACTP then to T30 and split by clinical response.

## Supplementary Information

**Supplementary Table 1.** Cell composition of ACT products based on scRNA-seq data.

**Supplementary Table 2.** List of annotated CD8^+^ T cell clonotypes. Tumor-reactive TCRs were validated by cloning and non-tumor-reactive ones were validated both by cloning and concomitant presence in scRNA-seq data.

**Supplementary Table 3.** Transcriptomic signature of CD8^+^ T cell tumor reactivity as computed by differential gene expression in single-cell RNA-seq data of baseline tumors (T0). The CD8^+^ T cell tumor reactivity signature was obtained by performing differential gene expression between CD8^+^ tumor-reactive (*n*=1499 cells) and non-tumor-reactive (*n*=1354 cells) T cells using linear regressions as inferred in the *lmFit* function of the *limma* R package. TCR-related genes and genes with average expression values lower than 0.3 were discarded from the differentially expressed gene list.

**Supplementary Table 4.** Transcriptomic changes of CD8^+^ T cell clonotypes tracked from baseline tumors (T0), to ACT products (ACTP), then to post-ACT tumor biopsies (T30). Differential Expressed Genes (DEG), transcription factors (DETF) and Reactome enrichment analyses in CD8^+^ T cells tracked from scRNA-seq data of T0 to ACTP, then to T30 are given.

